# Autophagy–Cholesterol Axis Remodeling Supports Malignant Progression and Chemoresistance in Glioma

**DOI:** 10.64898/2026.01.06.697885

**Authors:** Shahla Shojaei, Amir Barzegar Behrooz, Kianoosh Naghibzadeh, João Basso, Javad Alizadeh, Tania Dehesh, Roham Saberi, Bhavya Bhushan, Mehdi Eshraghi, Simone C da Silva Rosa, Courtney Clark, Mateusz Marek Tomczyk, Laura Cole, Grant Hatch, Vernon W Dolinsky, Vinith Yathindranath, Donald Miller, Christopher D. Pascoe, Sanjiv Dhingra, Abhay Srivastava, Amir Ravandi, Rui Vitorino, Stevan Pecic, Negar Azarpira, Zeinab Babaei, Mahmoud Aghaei, Saeid Ghavami

## Abstract

Glioma progression and resistance to temozolomide (TMZ) remain major clinical challenges. Here, we investigated whether dysregulated autophagy and cholesterol metabolism are coordinately remodeled during glioma progression and TMZ resistance. Tissue microarray analysis of astrocytoma and glioblastoma specimens revealed progressive autophagosome accumulation, reflected by increased LC3β puncta, coupled with impaired autophagic flux compared with adjacent normal brain tissue. These alterations intensified with tumor grade and were associated with upregulation of farnesyl diphosphate synthase (FDPS), linking malignant progression to cholesterol pathway remodeling.

TMZ-resistant (R) glioblastoma cells exhibited epithelial-to-mesenchymal transition, mitotic quiescence, and mitochondrial remodeling consistent with a therapy-tolerant phenotype. Bioenergetic profiling demonstrated reduced respiratory reserve, diminished ATP-linked respiration, and elevated proton leak, indicating constrained metabolic flexibility. In parallel, impaired autophagy flux was associated with suppression of de novo cholesterol synthesis and transcriptional downregulation of SREBP-2 and LDL-R.

Comprehensive lipidomic profiling revealed marked cholesterol metabolic reprogramming in R cells, characterized by accumulation of specific cholesteryl esters, including CE 22:5, CE 22:6, CE 22:4, and CE 20:4, despite reduced cholesterol biosynthesis. Pharmacologic inhibition of the mevalonate pathway with simvastatin significantly altered cholesteryl ester profiles but failed to restore autophagy flux or sensitize R cells to TMZ-induced apoptosis, even under combined TMZ–simvastatin treatment.

**Lay Abstract:** As gliomas progress from astrocytoma to glioblastoma, autophagy becomes dysregulated and cholesterol metabolism is rewired. This coordinated remodeling supports tumor survival, metabolic plasticity, and resistance to temozolomide therapy.

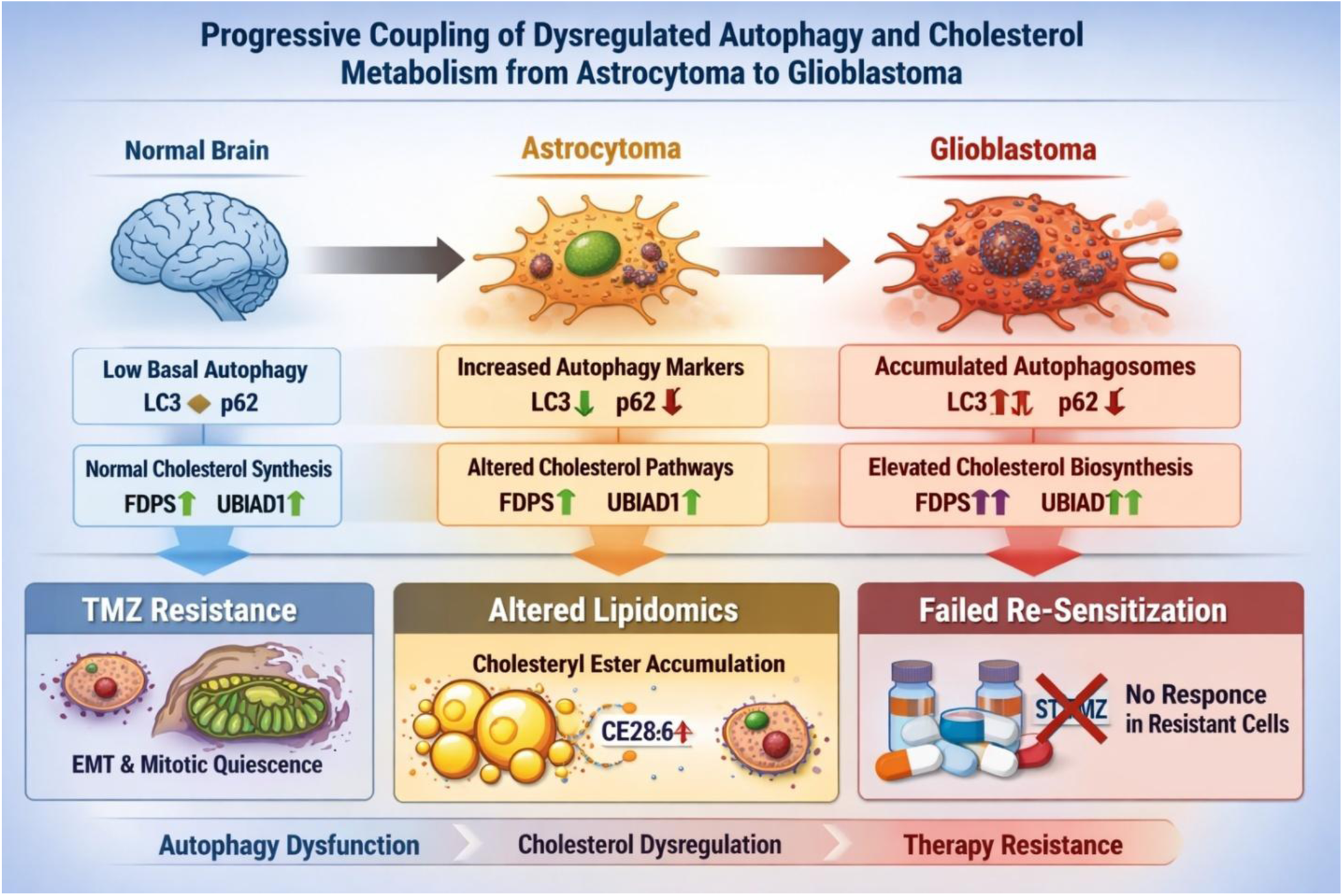

**Highlights:** Autophagy flux blockade intensifies during progression from astrocytoma to glioblastoma

Dysregulated autophagy is coupled to altered cholesterol metabolism in malignant gliomas

TMZ-resistant glioblastoma cells undergo epithelial-to-mesenchymal transition and mitotic quiescence

Resistant cells exhibit constrained bioenergetic capacity and mitochondrial remodeling

Impaired autophagy suppresses de novo cholesterol synthesis and lipid recycling

Lipidomics reveals accumulation of long-chain cholesteryl esters in TMZ-resistant cells

Statin-based cholesterol inhibition fails to resensitize glioblastoma cells to temozolomide

## Introduction

Glioblastoma (GB) stands as one of the most formidable and therapy-resistant brain tumors in adults. Conventional treatment strategies, comprising surgical excision followed by radiotherapy and temozolomide (TMZ)-based chemotherapy, offer limited effectiveness [1, 2]. The therapeutic efficacy of TMZ is frequently compromised by the onset of resistance, reducing its long-term impact[3, 4]. Emerging research has revealed that TMZ exposure can stimulate autophagy, a cellular survival mechanism crucial to GB’s resistance to treatment[5].

Autophagy serves as a paradoxical regulator in cancer, acting as both a guardian and a facilitator of tumor progression depending on cellular context [6]. This dynamic and multifaceted process influences various cancers, including glioblastoma, affecting their progression and therapeutic responses [7–9]. Autophagy dysfunction is a key driver of therapeutic resistance in GB, fueling both drug resistance and radio-resistance [10–12]. Beyond its role in treatment evasion, autophagy actively shapes GB pathophysiology by directly modulating cell survival, migration, and invasion while also rewiring metabolic networks to sustain tumor progression [12, 13].

GB cells exhibit metabolic adaptations that enhance their ability to proliferate rapidly and sustain their growth demands. One such adaptation involves reprogramming lipid metabolism, favoring de novo lipogenesis to support tumor progression [14, 15]. Recent findings indicate altered lipid homeostasis significantly drives resistance to TMZ in GB [16, 17]. Elevated intracellular cholesterol levels have been implicated in promoting TMZ resistance, highlighting the critical role of cholesterol and lipid metabolism in GB tumorigenesis [18, 19]. Villa et al. reported that, unlike astrocytes, GB cells do not rely on de novo cholesterol synthesis. Instead, they suppress key enzymes involved in cholesterol biosynthesis and predominantly acquire cholesterol from external sources[20]. Additional studies further support this observation, demonstrating that GB cells depend on low-density lipoprotein receptors (LDLR)-mediated cholesterol uptake, a process regulated by the EGFR/PI3K/Akt signaling axis [21, 22]. This metabolic shift underscores the rewiring of cholesterol metabolism in GB, enabling tumor cells to sustain their malignant behavior.

Research has demonstrated that disrupting lysosomal cholesterol efflux triggers autophagy-mediated suppression of GB cell proliferation. The opioid receptor agonists and antipsychotic drugs loperamide and pimozide drive autophagy-dependent cell death in MZ-54 GBM cells by engaging ATG5 and ATG7. These compounds hinder SMPD1, impair lysosomal activity and lead to ceramide accumulation, disrupting lysosomal degradation and its hexosyl derivatives. As a result, cholesterol becomes trapped within dysfunctional lysosomes, leading to oxidative stress-induced membrane destabilization. This process culminates in lysosomal membrane permeabilization (LMP), facilitating the release of cathepsin B (CTSB) into the cytosol, further promoting autophagy and cell death [23, 24]. The sterol regulatory element-binding protein 1 (SREBP-1) also plays a key role in regulating lipophagy to sustain cholesterol balance in GB cells. Activated in response to cholesterol scarcity, SREBP-1 enhances the expression of crucial autophagic genes such as ATG9B, ATG4A, and LC3B alongside the lysosomal cholesterol transporter NPC2. This activation fosters the degradation of lipid droplets (LDs) via lipophagy, facilitating the breakdown of cholesteryl esters (CEs) and the mobilization of cholesterol from lysosomes, ensuring the stability of plasma membrane cholesterol levels [25]. Thus, GB progression is closely linked to the interplay between cholesterol metabolism and autophagy.

Despite extensive work on autophagy and lipid metabolism in glioblastoma, it remains unclear whether dysregulation of these pathways arises as an early feature of gliomagenesis or progressively intensifies with malignant transformation. In particular, whether autophagy–cholesterol remodeling accumulates across tumor grade—from astrocytoma to glioblastoma—and predisposes tumors to stress tolerance and therapeutic resistance has not been systematically examined. Moreover, while autophagy is frequently described as “activated” in GB [26–28], increasing evidence suggests that high-grade tumors may instead exhibit a state of autophagy flux blockade, characterized by autophagosome accumulation without efficient lysosomal degradation. Such a state could fundamentally alter lipid handling and metabolic feedback regulation.

Importantly, impaired autophagy flux is predicted to disrupt endogenous lipid recycling [29–31], raising the possibility that metabolic adaptations in resistant glioblastoma cells may not be reversible by simple replenishment or inhibition of cholesterol biosynthesis alone. Whether pharmacologic modulation of cholesterol availability—through exogenous lipid delivery or HMG-CoA reductase inhibition—can overcome resistance in the context of autophagy flux blockade remains unresolved. Addressing this question is essential, as it challenges prevailing assumptions that correcting metabolic deficits is sufficient to restore chemosensitivity.

In this study, we systematically dissect the relationship between autophagy dysregulation, cholesterol metabolism, and metabolic reprogramming across glioma progression and in the setting of temozolomide resistance. Using patient-derived tissue microarrays, resistant glioblastoma models, lipidomics, and mitochondrial functional analyses, we demonstrate that autophagy flux blockade and cholesterol pathway remodeling intensify with tumor grade and stabilize a stress-tolerant, therapy-resistant phenotype. We further show that pharmacologic modulation of cholesterol metabolism fails to restore chemosensitivity once this adaptive program is established. Together, these findings define an autophagy–cholesterol axis as a central organizing principle of malignant progression and chemoresistance in glioblastoma.

## MATERIALS AND METHODS

### Cell Culture

All experiments were performed using the U251-mKate human glioblastoma cell line, originally developed by Dr. Marcel Bally (Experimental Therapeutics, British Columbia Cancer Agency, Vancouver, BC, Canada). Cells were maintained in high-glucose Dulbecco’s modified Eagle’s medium (DMEM; 4 mg/mL glucose; Corning, 50-003-PB) supplemented with 10% fetal bovine serum (FBS; Gibco™, 16000044) and 1% penicillin–streptomycin. Cultures were maintained under standard humidified conditions at 37 °C with 5% CO₂ and routinely monitored to ensure exponential growth and absence of contamination prior to experimentation.

### Immunohistochemistry for Autophagy and Cholesterol Metabolism Markers

Tissue microarray (TMA) sections containing astrocytoma, glioblastoma, and matched adjacent non-neoplastic brain tissues were used for immunohistochemical analysis of autophagy and cholesterol metabolism–associated proteins. Formalin-fixed, paraffin-embedded tissue sections (4 μm thickness) were deparaffinized in xylene and rehydrated through a graded ethanol series to distilled water.

Antigen retrieval was performed by heating sections in citrate buffer (10 mM sodium citrate, pH 6.0) using a pressure cooker or microwave for 15–20 min, followed by gradual cooling to room temperature. Endogenous peroxidase activity was quenched by incubating sections in 3% hydrogen peroxide for 10 min. Non-specific binding was blocked by incubation with 5% normal serum (matched to the secondary antibody host species) for 30 min at room temperature.

Sections were incubated overnight at 4 °C with the following primary antibodies diluted in antibody diluent: anti-LC3β (for autophagosome detection), anti-p62/SQSTM1 (for assessment of autophagic flux), anti-FDPS (ab153805, Abcam; 1:200 dilution), and anti-UBIAD1 (NBP2-82046, Novus Biologicals; 1:200 dilution). Each marker was stained on serial sections under identical conditions.

After washing in phosphate-buffered saline (PBS), sections were incubated with appropriate HRP-conjugated secondary antibodies for 30 min at room temperature. Immunoreactivity was visualized using 3,3′-diaminobenzidine (DAB) as chromogen, followed by counterstaining with hematoxylin. Sections were dehydrated, cleared, and mounted using permanent mounting medium.

Immunostained slides were digitally scanned or examined under light microscopy. Cytosolic puncta formation for LC3β and p62 was evaluated as a surrogate for autophagosome abundance and autophagic flux, respectively. FDPS and UBIAD1 expression were assessed based on cytoplasmic staining intensity. Staining intensity and puncta density were semi-quantitatively scored by at least two independent observers blinded to clinicopathological data. Discrepancies were resolved by joint review. Quantitative and categorical scoring results were used for downstream statistical and correlation analyses.

### Generation of a Resistant GBM Cell Line

U251-mKate cells were cultured in T75 tissue culture flasks in a complete medium containing high glucose DMEM, 10% FBS (Gibco, 10437-036), and 1% penicillin-streptomycin (Gibco, 15140122) and incubated at 37°C, 5% CO_2_, in a humidified incubator (Thermo Fischer Scientific, USA). A pulsed-selection strategy was chosen to make a clinically relevant chemoresistant model [32]. Once cells reached 70% confluence, they were treated with 100 μM TMZ (Sigma Aldrich, T2577) over three weeks, followed by four weeks of TMZ-free medium for recovery. Cells were cultured in complete medium with 250 μM TMZ for three weeks, followed by another four weeks with TMZ-free medium. The final population of U251-mKate cells resistant to 250 μM TMZ was selected and cultured in a complete medium without TMZ for four weeks.

### Immunoblotting

Protein expression was analyzed by Western blotting, using GAPDH and Ponceau S staining as loading and transfer controls. Cells were lysed in ice-cold lysis buffer containing 20 mM Tris–HCl (pH 7.5), 0.5% Nonidet P-40, 0.5 mM phenylmethylsulfonyl fluoride (PMSF; Sigma-Aldrich, P7626), 100 μM β-glycerol 3-phosphate (Sigma-Aldrich, 50020), and 0.5% protease and phosphatase inhibitor cocktail (Sigma-Aldrich, PPC1010). Lysates were clarified by centrifugation at 10,000 × g for 10 min at 4 °C, and supernatants were collected for analysis. Protein concentrations were determined using the Lowry assay. Equal amounts of protein were resolved by SDS–polyacrylamide gel electrophoresis and transferred onto polyvinylidene difluoride (PVDF) membranes. Membranes were blocked with 5% skim milk in Tris-buffered saline containing 0.1% Tween-20 (TBST) for 1 h at room temperature, followed by incubation with primary antibodies diluted in 1% skim milk in TBST overnight at 4 °C. After washing, membranes were incubated with horseradish peroxidase (HRP)–conjugated secondary antibodies for 2 h at room temperature. Immunoreactive bands were detected using enhanced chemiluminescence (ECL; Amersham-Pharmacia Biotech) and visualized using a Bio-Rad imaging system. Band intensities were quantified by densitometric analysis using AlphaEase FC software [33, 34].

### Transmission Electron Microscopy (TEM)

Autophagic structures and mitochondrial ultrastructure were examined using transmission electron microscopy (TEM). Cells were cultured in 100-mm dishes in Dulbecco’s modified Eagle’s medium (DMEM) supplemented with 10% fetal bovine serum and 1% penicillin–streptomycin under standard humidified incubator conditions (37 °C, 5% CO₂). Cells were harvested and immediately fixed in Karnovsky’s fixative (Sigma-Aldrich, SBR00124) for 1 h at 4 °C. Following fixation, cell pellets were resuspended in 5% sucrose prepared in 0.1 M Sørensen’s phosphate buffer and incubated overnight at 4 °C. Samples were then pelleted and post-fixed with 1% osmium tetroxide in 0.1 M Sørensen’s phosphate buffer for 2 h at room temperature. Cells were subsequently dehydrated through a graded ethanol series and embedded in Embed 812 resin (Sigma-Aldrich, 45347). Ultrathin sections (∼200 nm) were cut using an ultramicrotome and mounted on copper grids. Sections were stained with uranyl acetate followed by lead nitrate for contrast enhancement. Ultrastructural imaging was performed using a Philips CM10 transmission electron microscope, allowing detailed visualization of autophagosomes, mitochondrial morphology, and intracellular membrane organization [34].

### Cell Proliferation by Trypan Blue Exclusion and Direct Cell Counting

Baseline proliferation of U251 glioblastoma NR and R cells was quantified by direct cell counting using Trypan Blue exclusion. Cells were maintained in high-glucose DMEM supplemented with 10% FBS and 1% penicillin/streptomycin. To ensure comparable starting confluence between groups, NR and R cells were seeded at equal densities (50,000 cells/mL) in tissue-culture plates and allowed to grow under standard conditions (37°C, 5% CO₂).

At the indicated time points (72h), both floating and adherent cells were collected. Culture supernatants were retained, adherent cells were detached using trypsin–EDTA, and the fractions were combined to avoid underestimation of cell loss. Cell suspensions were mixed 1:1 with 0.4% Trypan Blue, and viable (Trypan Blue–negative) and non-viable (Trypan Blue–positive) cells were counted using a hemocytometer (or an automated cell counter). Proliferation was expressed as the total number of viable cells per well at each time point. Viability (%) was calculated as: (viable cells / total cells) × 100, and fold-change in viable cell number relative to baseline (time 0) was reported where indicated.

### Flow Cytometry/Cell Cycle Analysis

U251 glioblastoma cells were seeded in 12-well plates at a density of 50,000 cells/well for TMZ-non-resistant (NR) cells and 75,000 cells/well for TMZ-resistant (R) cells and allowed to grow to approximately 60% confluence. Cells were then treated with temozolomide (TMZ; 250 µM) for 48 h. Following treatment, cells were detached using EDTA buffer, collected by centrifugation at 300 × g for 10 min, and washed once with phosphate-buffered saline (PBS). Cell pellets were resuspended in hypotonic propidium iodide (PI) lysis buffer containing 0.01% Triton X-100, 1% sodium citrate, and 40 µg/mL PI, and incubated for 30 min at room temperature in the dark.

DNA content was analyzed by flow cytometry using a Beckman Coulter CytoFLEX LX flow cytometer equipped with a 488-nm laser and a 610/20 band-pass filter. Cell populations corresponding to sub-G0/G1 (apoptotic), G0/G1, S, and G2/M phases were quantified based on DNA content [35].

### Measurement of Mitochondrial Respiration

Mitochondrial respiration was assessed by measuring the oxygen consumption rate (OCR) using an Agilent Seahorse XFe24 extracellular flux analyzer. NR and R glioblastoma cells were seeded at a density of approximately 0.3 × 10⁶ cells per well in XF24 cell culture microplates in complete growth medium and allowed to adhere overnight. On the day of the assay, cells were washed twice and the medium was replaced with Seahorse XF Base Minimal DMEM supplemented with glucose, L-glutamine, and sodium pyruvate (Agilent Technologies, #103334-100). Cells were then incubated at 37 °C in a non-CO₂ incubator for 1 h prior to analysis. Baseline OCR was recorded, followed by sequential injections of mitochondrial modulators to interrogate respiratory parameters. Proton leak–associated respiration was measured following the injection of oligomycin (1 µM). Maximal respiratory capacity was determined by addition of the mitochondrial uncoupler FCCP (2 µM). Non-mitochondrial respiration was assessed after combined injection of rotenone (1 µM) and antimycin A (1 µM). OCR data were normalized to live cell number by staining individual wells with Hoechst 33342 (Invitrogen, #R37605) and quantifying nuclear signal using a Cytation 5 imaging system (BioTek). Maximal respiration was calculated by subtracting non-mitochondrial OCR from FCCP-stimulated OCR. Spare respiratory capacity was determined by subtracting baseline OCR from FCCP-stimulated OCR. ATP-linked respiration was calculated by subtracting oligomycin-treated OCR from baseline OCR, and proton leak was calculated by subtracting non-mitochondrial OCR from oligomycin-treated OCR. Data are presented as pmol O₂ per minute per cell.

### Gene Expression Analysis

Total RNA was extracted according to the manufacturer’s instructions using the PureLink RNA Mini Kit (Invitrogen, #12183018A). RNA concentration and purity were assessed, and 500 ng of total RNA was reverse-transcribed into complementary DNA (cDNA) using the LunaScript RT SuperMix Kit (New England Biolabs, #E3010L). Quantitative real-time PCR (qPCR) was performed using the Luna Universal qPCR Master Mix (New England Biolabs, #M3003X) with 10 ng cDNA per well in a 96-well plate format on a CFX96 Real-Time PCR Detection System (Bio-Rad).

Gene expression of LDL receptor (LDL-R) and sterol regulatory element-binding protein 2 (SREBP-2) was quantified using the following primers: LDL-R forward, 5′-CCCGACCCCTACCCACTT-3′; LDL-R reverse, 5′-AATAACACAAATGCCAAATGTACACA-3′; SREBP-2 forward, 5′-GACGCCAAGATGCACAAGTC-3′; SREBP-2 reverse, 5′-ACCAGACTGCCTAGGTCGAT-3′.

For normalization, β-ACTIN (forward, 5′-CACCATTGGCAATGAGCGGTT-3′; reverse, 5′-AGGTCTTTGCGGATGTCCACGT-3′), EIF2α (forward, 5′-CGAAACACTGTCTCTCAGTCAA-3′; reverse, 5′-CCAGTTGCTGCTTGTTCTTTC-3′), and 18S rRNA (forward, 5′-GCAGAATCCACGCCAGTACAAG-3′; reverse, 5′-GCTTGTTGTCCAGACCATTGGC-3′) were used as reference genes. Gene expression levels were normalized using the geometric mean of β-ACTIN, EIF2α, and 18S.

Primer specificity was verified by in silico PCR using the ENCODE UCSC Genome Browser and experimentally confirmed by electrophoresis of PCR products on a 1% agarose gel prepared in 1× Tris-Acetate-EDTA (TAE) buffer (MP Biomedicals, #TAW50X01) at 100 V for 45 min.

### MitoView Live-Cell Imaging

Live-cell mitochondrial imaging was performed using MitoView™ Green (Biotium, #70054) to assess mitochondrial morphology and network organization, and Hoechst 33342 (Biotium, #33342) to visualize cell nuclei. NR and R glioblastoma cells were cultured on glass-bottom dishes (or tissue-culture–treated plates) under standard growth conditions. Cells were incubated with MitoView Green at a final concentration of 100 nM and Hoechst 33342 at 5 µM, prepared in pre-warmed complete culture medium. Staining was performed for 15–30 min at 37 °C in a humidified incubator protected from light. Following incubation, cells were gently washed with pre-warmed phosphate-buffered saline to remove excess dye and maintained in fresh, phenol red–free medium for imaging. Live-cell fluorescence imaging was carried out immediately using an epifluorescence or confocal microscope equipped with appropriate excitation/emission filters. Imaging parameters were kept constant across all experimental conditions to allow direct comparison of mitochondrial morphology between NR and R cells.

### Preparation and Characterization of Cholesterol-Containing Lipid Nanoparticles (LNPs)

Cholesterol-containing lipid nanoparticles (LNPs) were prepared using a microfluidic mixing approach. Lipid components, including 1,2-dioleoyl-sn-glycero-3-phosphocholine (DOPC), 1,2-distearoyl-sn-glycero-3-phosphocholine (DSPC), cholesterol, 1,2-dimyristoyl-rac-glycero-3-methoxypolyethylene glycol-2000 (DMG-PEG), and the fluorescent lipid tracer DiO, were obtained from Avanti Polar Lipids/Sigma-Aldrich (St. Louis, MO, USA). Lipids were dissolved in ethanol at a molar ratio (%) of 50:10:38:1.5:0.5 (DOPC:DSPC:cholesterol:DMG-PEG:DiO), and the total lipid concentration of the organic phase was maintained at 10 mg/mL.

LNPs were formed by rapidly mixing 0.5 mL of the ethanol-based lipid solution with 1.5 mL of aqueous phosphate-buffered saline (PBS, pH 7.4) using a herringbone-patterned microfluidic cartridge on a NanoAssemblr Benchtop instrument (Precision NanoSystems, Vancouver, BC, Canada). Microfluidic assembly was performed at a flow rate ratio (FRR) of 1:3 (organic phase:aqueous phase) and a total flow rate (TFR) of 12 mL/min, enabling controlled and reproducible nanoparticle self-assembly.

Following formulation, LNP suspensions were purified to remove residual ethanol by centrifugal filtration using a 10 kDa molecular weight cut-off membrane at 2000 × g with PBS (pH 7.4). Purified LNPs were concentrated to a final volume of 0.5 mL, corresponding to a lipid concentration of 10 mg/mL, and stored at 4 °C until use.

The hydrodynamic diameter, polydispersity index (PDI), and surface charge (zeta potential) of the LNPs were determined by dynamic light scattering using a ZetaPALS analyzer (Brookhaven Instruments, NY, USA). For measurements, LNPs were diluted in PBS (pH 7.4) to a final lipid concentration of 20 µg/mL. The formulated LNPs exhibited a mean hydrodynamic diameter of 65.3 ± 1.4 nm, a near-neutral zeta potential (−0.68 ± 2.3 mV), and a PDI of 0.151 ± 0.038, indicating a stable and monodisperse nanoparticle population under physiological conditions.

### Phase Contrast

Bright-field images were acquired using an Axio Observer microscope (ZEISS, Oberkochen, Germany) equipped with 20× and 50× objective lenses. Image processing, including background subtraction, scale bar calibration, and quantitative analysis, was performed using Fiji (ImageJ) and ZEN 2.3 Pro software (ZEISS).

### Cholesterol Mass Measurement

NR and R glioblastoma cells were cultured in cholesterol-free medium and harvested at 24, 48, and 72 h. Cellular cholesterol content was quantified according to the manufacturer’s instructions using the Amplex Red Cholesterol Assay Kit (Invitrogen, Thermo Fisher Scientific, Toronto, Canada), as previously described [36, 37]. Cholesterol levels were normalized to total cellular protein content.

### De Novo Cholesterol Biosynthesis and Cholesterol Esterification

De novo cholesterol biosynthesis and cholesterol esterification were assessed using a modified Mokashi protocol [37]. Briefly, NR and R cells were cultured in cholesterol-free medium supplemented with ITS (insulin–transferrin–selenium) for 24, 48, and 72 h. Cells were then incubated in serum-free DMEM containing 2 µCi [¹⁴C]-acetate per 2 mL dish for the indicated time periods. Following incubation, cells were harvested and washed twice with ice-cold PBS (pH 7.4). Total lipids were extracted as described previously [38], and incorporation of radiolabeled acetate into free cholesterol and cholesteryl esters was quantified and normalized to total protein content, as previously reported [39].

### MTT Cell Viability Assay

Cell viability was assessed using the MTT assay as previously described [35], with modifications to evaluate the effects of LNP and TMZ in NR and R U251 glioblastoma cells, as well as in primary astrocytes.

Cells were cultured in high-glucose DMEM supplemented with 10% fetal bovine serum and 1% penicillin/streptomycin and seeded into 96-well plates (Corning, #353072). To achieve comparable baseline confluence, NR cells were seeded at 2,300 cells/well and R cells at 3,100 cells/well in a final volume of 200 µL per well. Cells were allowed to adhere overnight and treated once cultures reached approximately 50% confluence. Cells were treated with LNP (5, 10, 25, or 100 µg/mL), TMZ (100 or 200 µM), or their combinations for 72 h. Control wells received vehicle only. All treatments were prepared in complete DMEM containing 10% FBS. Primary astrocytes were treated in parallel to assess CE-NP toxicity in non-tumor cells. Following treatment, MTT reagent (20 µL of 5 mg/mL stock solution in PBS; final concentration 0.5 mg/mL) was added directly to each well and plates were incubated for 3 h at 37 °C in 5% CO₂, protected from light. The culture medium was then carefully aspirated, and formazan crystals were solubilized by adding 200 µL dimethyl sulfoxide (DMSO) per well with gentle mixing. Absorbance was measured immediately at 570 nm using a Synergy H1 microplate reader (BioTek). Background absorbance from media-only wells was subtracted. Cell viability was calculated as a percentage relative to untreated control cells using the formula: (OD₅₇₀ treated / OD₅₇₀ control) × 100. Data were analyzed using Prism software, and statistical significance was determined by two-way ANOVA.

### Liquid Chromatography-Mass Spectrometry (LC-MS) for Lipidomics Analysis of Cholesterol Species

Lipids were extracted from U251 GB cells (NR, R, NR-BCL2L13-KD, R-BCL2L13-KD) using a chloroform:methanol (2:1, v:v) mixture. Cells grown to confluence were collected, sonicated, and centrifuged (1000 g, 10 min). The supernatant was mixed with the internal standard (ISTD), vortexed, and centrifuged (3500 rpm, 5 min). The lower phase was dried under N₂ gas and reconstituted in water-saturated butanol, followed by methanol with 10 mM ammonium formate. After centrifugation (10,000g, 10 min), lipids were analyzed by LC-MS. Separation was performed on a Zorbax C18 column using a linear gradient of mobile phases A and B, both containing 10 mM ammonium formate. The elution program involved increasing from 0% to 100% mobile phase B over 8 min, with re-equilibration to starting conditions before the next injection. Diacylglycerol and triacylglycerol species were separated isocratically at 100 μl/min. The column was maintained at 50°C, and the injection volume was 5 μl.

Lipids eluted from the HPLC system were introduced into the AbSciex 4000 QTRAP triple quadrupole linear hybrid mass spectrometer. The mass spectrometer was operated in scheduled Multiple Reaction Monitoring (MRM) mode. Unique lipids (322) spanning 25 lipid classes/subclasses were screened for targeted semi-quantification. All lipid species other than fatty acids were scanned in positive electrospray ionization mode [ESI+]. The individual lipids in each lipid class were identified by lipid class-specific precursor ions or neutral losses. Lipids were then quantified by comparing the deisotoped lipid peak areas against those of the class-specific ISTDs added before lipid extraction. The total carbon number of the fatty acids represented lipids. In ESI+ mode, the instrument settings were optimized as follows: curtain gas (psi), 26; collision gas (nitrogen), medium; ion spray voltage (V), 5500; temperature (°C), 500.0; ion source gas 1 (psi), 40.0; ion and source gas 2 (psi), 30.0. The MRM detection window was fixed between 45 s and 90 s depending upon the chromatographic peak width of the lipid class. Isobaric species within the same class, such as PC(O) and PC(P), exhibited clear separation in this method. Also, molecular species within the same lipid class, which differ only in the number of double bonds, were well separated chromatographically. All analyses were performed with R software (version 4.1.1). Limma, pheatmap, and ggplot2 packages were used.

### Lipidomics Analysis

Cholesterol-related lipidomic data were normalized and visualized using R software (version 4.1.1) and MetaboAnalyst 6.0 to evaluate treatment-dependent alterations in cholesterol profiles and to assess differences between NR and R conditions. Multivariate statistical analysis was performed using partial least squares discriminant analysis (PLS-DA) to identify key cholesterol-associated lipids contributing to the discrimination between NR and R samples. PLS-DA was applied both across all treatment and control groups collectively, as well as separately for each treatment condition. Lipids with a variable importance in projection (VIP) score > 1.4 were considered as a threshold for key contributor lipids to group discrimination and were retained for downstream analysis (**detailed results and analyses are provided in supplementary**).

To investigate the biological pathways potentially modulated by the identified key contributor lipids, the Human Metabolome Database (HMDB; https://hmdb.ca) was used to retrieve the corresponding SMILES codes. These SMILES representations were subsequently submitted to SwissTargetPrediction (https://www.swisstargetprediction.ch/) to predict putative protein targets associated with each lipid. The resulting protein targets were annotated using their UniProt identifiers and subjected to protein–protein interaction and functional enrichment analysis using the STRING (Version 12.0; https://string-db.org/) database. Based on these interactions and associated gene sets, KEGG pathway enrichment analysis was performed to identify signaling pathways potentially influenced by the altered cholesterol lipid profiles.

### Statistics and reproducibility

All experiments were conducted in triplicate, with three independent repeats. Data are presented as mean ± standard deviation. Statistical significance was determined using a two-tailed t-test or one-way/two-way ANOVA with Tukey’s post hoc test for multiple comparisons, considering p-values ≤ 0.05 as significant. Lipid classification and annotation were performed using the LIPIDMAPS database. Lipidomic data were analyzed through PLS-DA test by considering VIP score > 1.4 as significant. Associations between categorical clinicopathological variables (tumor type, age group, gender, and WHO grade) and protein expression levels (cytosolic LC3, P62, FDPS, and UBIAD) as well as autophagic flux categories were evaluated using Fisher’s exact test for contingency tables with small sample sizes. Pearson’s χ² test was applied where cell counts met the assumptions for χ² analysis. Data are presented as percentages within each category. All statistical tests were two-sided, and a p value < 0.05 was considered statistically significant. Missing values (NA) indicate insufficient data for correlation estimation in specific subgroups.

## Results

### Progressive Coupling of Dysregulated Autophagy and Cholesterol Metabolism from Astrocytoma to Glioblastoma

To investigate whether autophagy and cholesterol metabolism are coordinately altered during glioma progression, we performed a comprehensive tissue microarray–based analysis of autophagy-associated markers (LC3β and p62/SQSTM1) and cholesterol/isoprenoid biosynthesis enzymes (FDPS and UBIAD1) in astrocytoma and glioblastoma specimens, together with matched adjacent non-neoplastic brain tissues.

Autophagosome-associated LC3β was assessed by quantification of cytosolic LC3 puncta, which reflect steady-state autophagosome abundance [23, 36, 40–42]. Adjacent normal brain tissues exhibited low LC3 puncta density, consistent with basal autophagy. In contrast, both astrocytoma and glioblastoma specimens demonstrated significantly increased LC3β puncta, with glioblastoma displaying a markedly higher puncta burden than astrocytoma, indicating subtype- and grade-dependent autophagosome accumulation (**Figure 1A**, **Table 1**). Overall, strong LC3β puncta expression was observed in 63 of 68 tumor cores (92.6%), with 67 of 68 tumors (98.5%) showing LC3β positivity, whereas normal tissues showed predominantly weak or moderate puncta intensity (P < 0.0001; **Table 1**).

**Figure 1:**
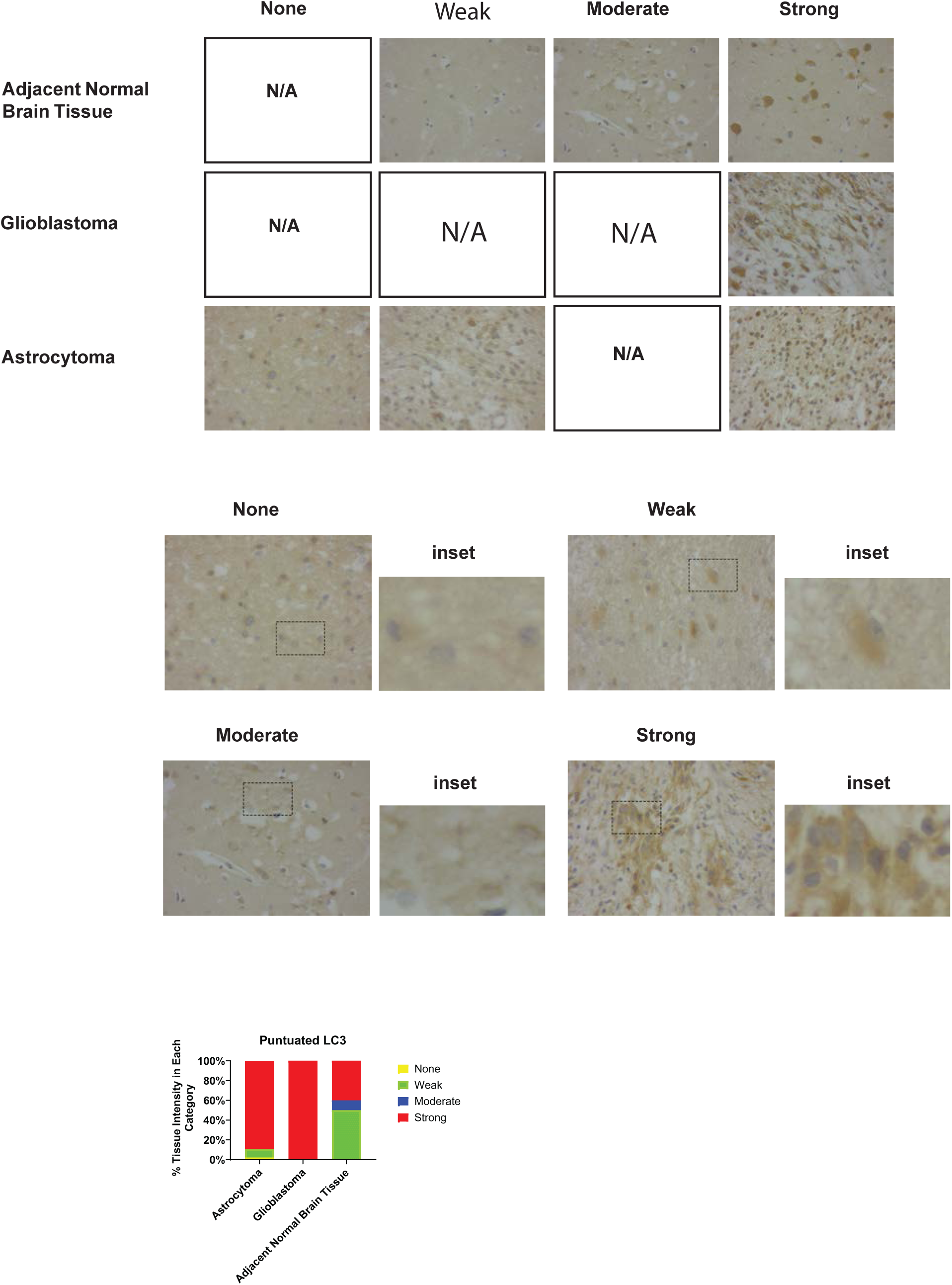
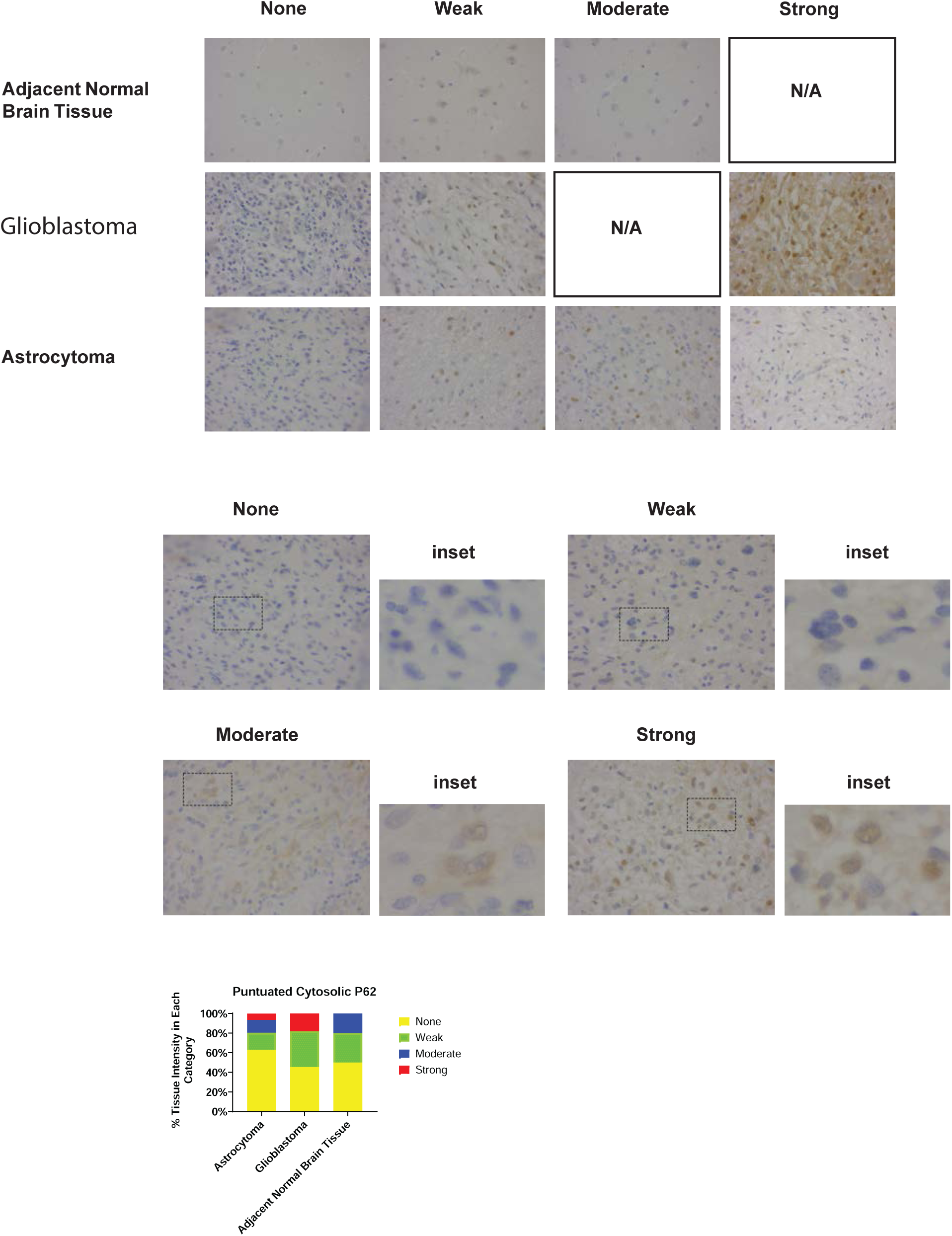
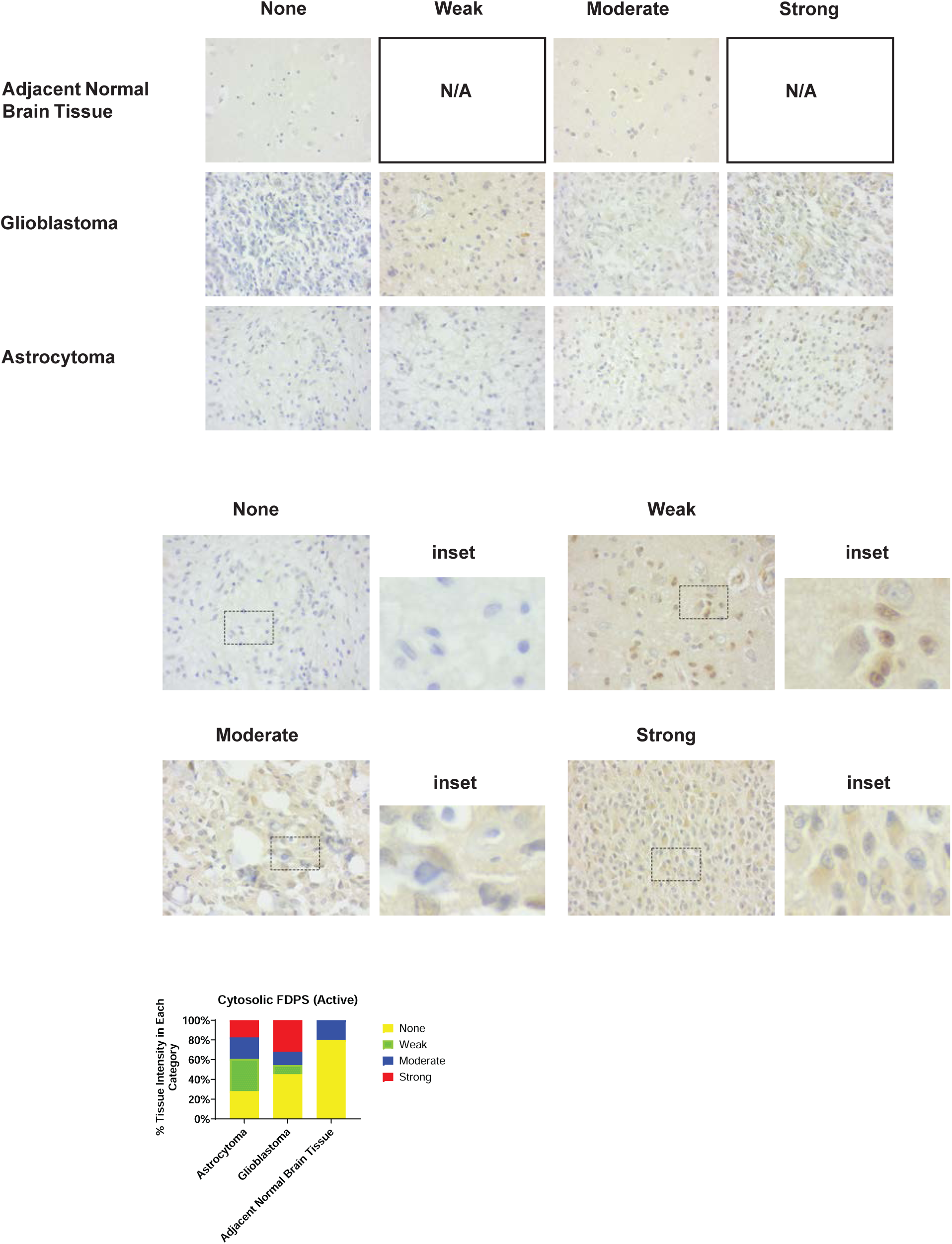
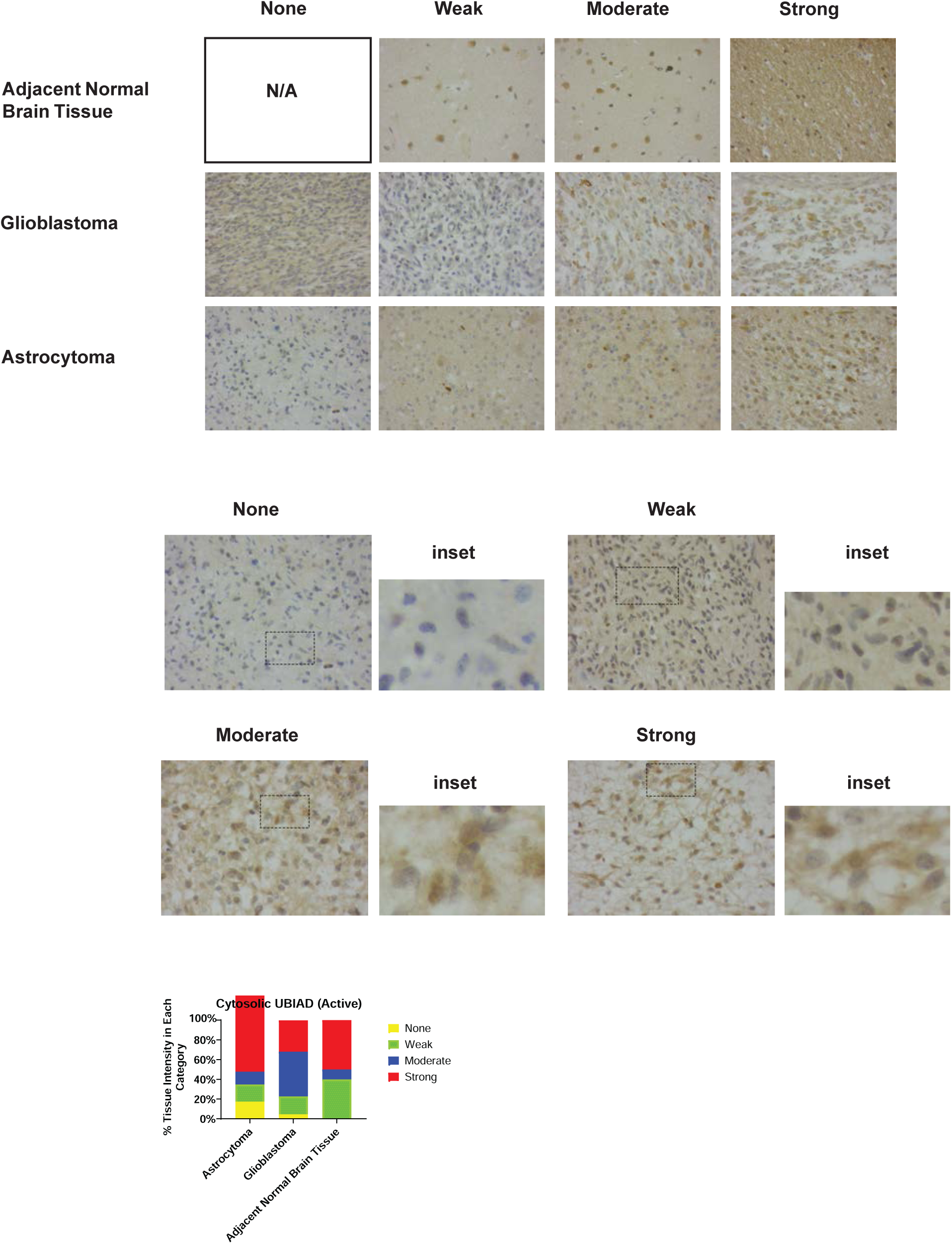
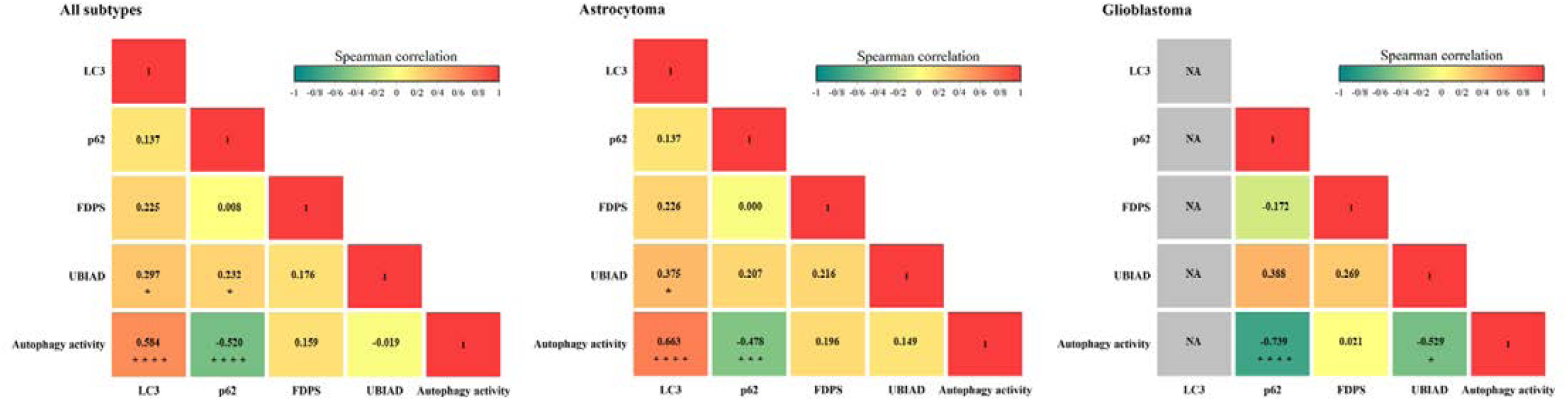
Immunohistochemical Staining for Autophagy and Cholesterol Metabolism Proteins in Human Brain Tissues. (A) LC3 puncta Expression: LC3 puncta, an autophagosome marker, was evaluated in brain tissue using TMA. Three pathologists assessed the staining blindly, categorizing it as None, Weak, Moderate, or Strong. Results showed significantly higher LC3 puncta in astrocytoma and GB tissues compared to normal brain tissue (P < 0.0001). LC3 expression is linked to astrocytoma (P < 0.01) and GB (P < 0.0001) tumors. Scale bar = 50 µM. (B) p62 Expression: p62, a marker for autophagosome degradation, was similarly evaluated. No significant difference in p62 expression was found between normal and tumor tissues (P = 0.499). p62 expression is not associated with astrocytoma (P = 0.644) or GB (P = 0.128) tumors. Scale bar = 50 µM. (C) FDPS Expression: FDPS, a cholesterol biosynthesis pathway enzyme, in its active form, was detected in brain tissue TMA. Results indicated significant differences in FDPS expression between normal and tumor tissues (P < 0.01), with associations found for astrocytoma (P < 0.5) and GB (P < 0.01). Scale bar = 50 µM. (D) UBIAD Expression:UBIAD, a cholesterol biosynthesis pathway enzyme, also in its active form, was analyzed with no significant difference in expression between normal and tumor tissues (P = 0.344). UBIAD is not associated with astrocytoma (P = 0.344) or GB (P = 0.213) tumors. Scale bar = 50 µM. (E) Correlation Analysis: Spearman’s correlation analysis examined the relationship between autophagy and MVA pathway proteins in different brain tumors. Significant correlations are marked as * (P ≤ 0.05), *** (P ≤ 0.001), and **** (P ≤ 0.0001).

**Table 1.**
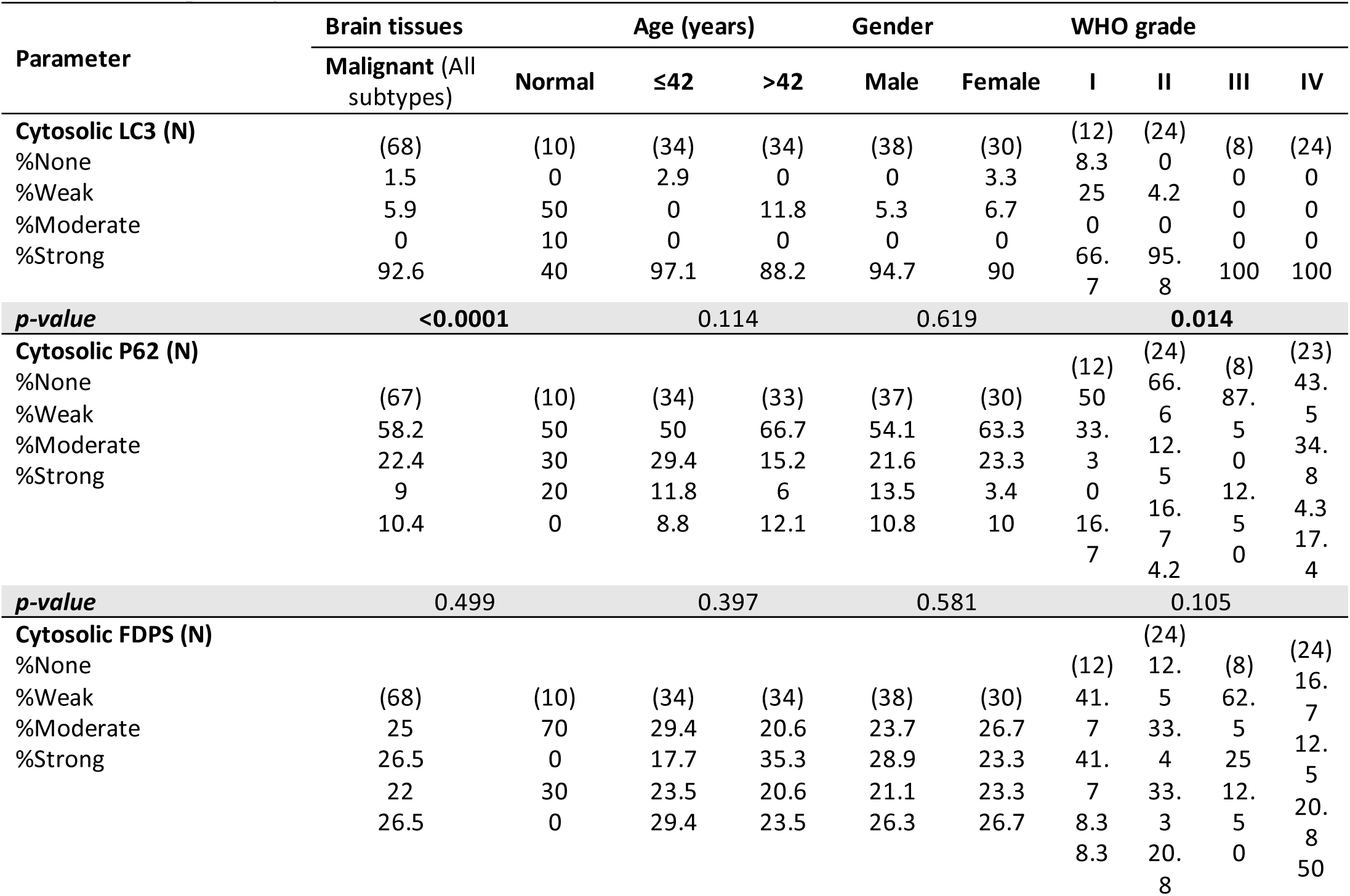
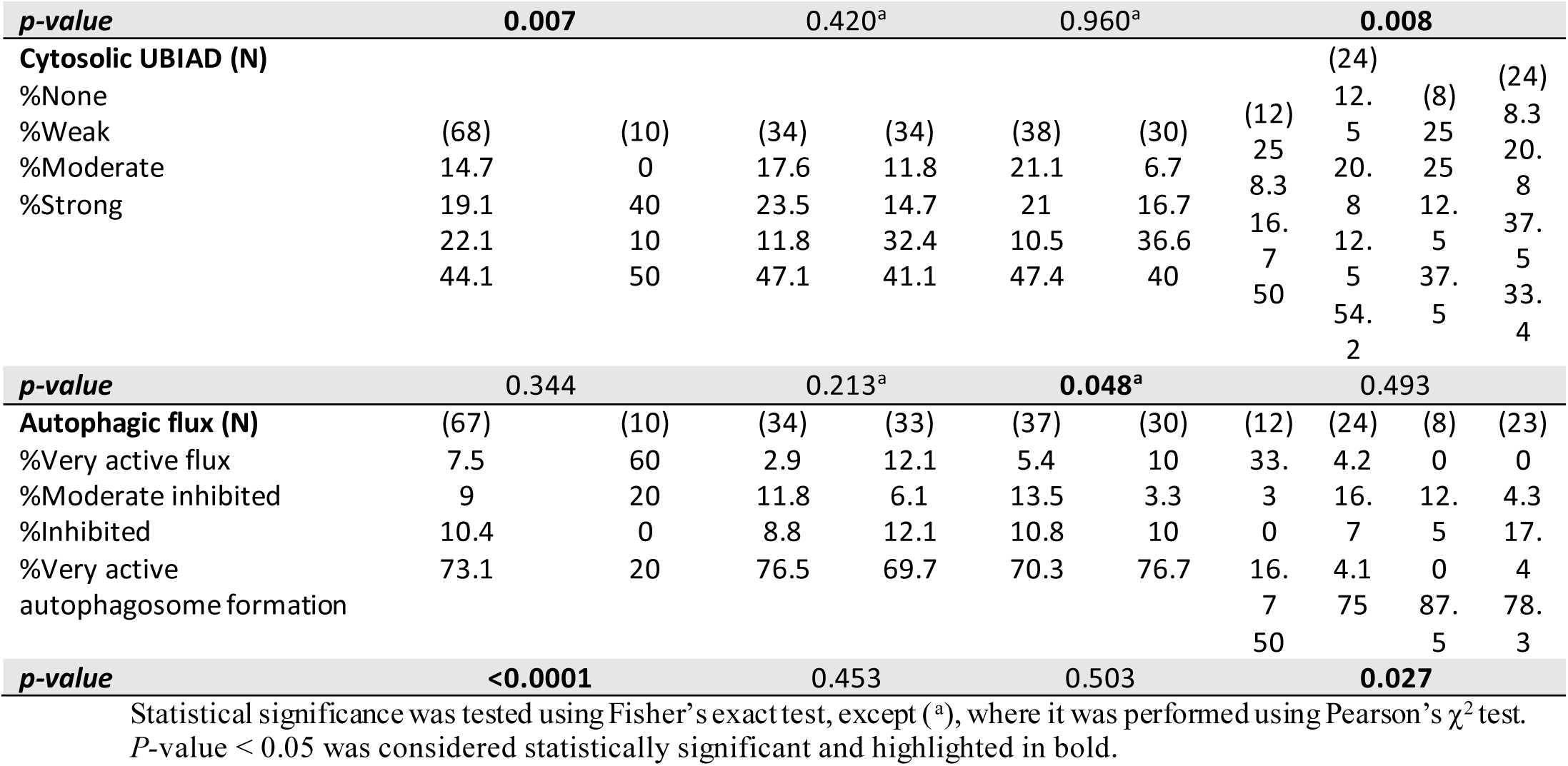
Correlation of cytosolic LC3, P62, FDPS, and UBIAD protein expression and autophagic flux with clinicopathological features of Brain tumor

We next evaluated p62/SQSTM1, a selective autophagy adaptor that binds LC3β and is degraded during effective autophagic flux [3]. Immunohistochemical analysis revealed absent or weak p62 puncta in the majority of tumor specimens (54 of 67; 80.6%), with no significant difference in overall p62 positivity between tumor and normal tissues (P = 0.499; **Figure 1B**, **Table 1**). Integration of LC3β and p62 puncta using established criteria allowed inference of relative autophagy states [42]. Among tumor tissues, the predominant pattern was autophagosome accumulation (49 of 67; 73.1%), whereas only a small fraction (5 of 67; 7.5%) exhibited features consistent with relatively preserved autophagic flux (**Table 1**). In contrast, normal brain tissues more frequently demonstrated effective autophagic flux (6 of 10; 60%), confirming a tumor-associated shift toward autophagosome accumulation (P < 0.0001; **Table 1**).

Given the emerging links between autophagy and lipid metabolism in cancer, we examined the expression of FDPS and UBIAD1, two key enzymes involved in cholesterol and isoprenoid biosynthesis [43–45]. FDPS expression was significantly elevated in tumor tissues (51 of 68; 75.0%) compared with normal brain tissues (3 of 10; 30.0%) (P = 0.007; **Figure 1C**, **Table 1**), implicating FDPS upregulation in glioma-associated metabolic reprogramming. In contrast, UBIAD1 expression was detected in most tumors (58 of 68; 85.3%) but did not differ significantly between tumor and normal tissues (P = 0.344; **Figure 1D**, **Table 1**).

Subtype-specific analyses further demonstrated that astrocytoma and glioblastoma differ in the magnitude of these alterations. Astrocytomas showed significantly increased LC3β puncta, FDPS expression, and autophagosome accumulation relative to normal tissues, but retained greater heterogeneity in autophagy states (**Table 2**). Glioblastomas, by contrast, uniformly exhibited strong LC3β puncta and were overwhelmingly classified as autophagosome-accumulating, with significantly higher FDPS expression than normal tissues (**Figure 1A–D**, **Table 3**). UBIAD1 expression remained largely independent of tumor subtype.

**Table 2.**
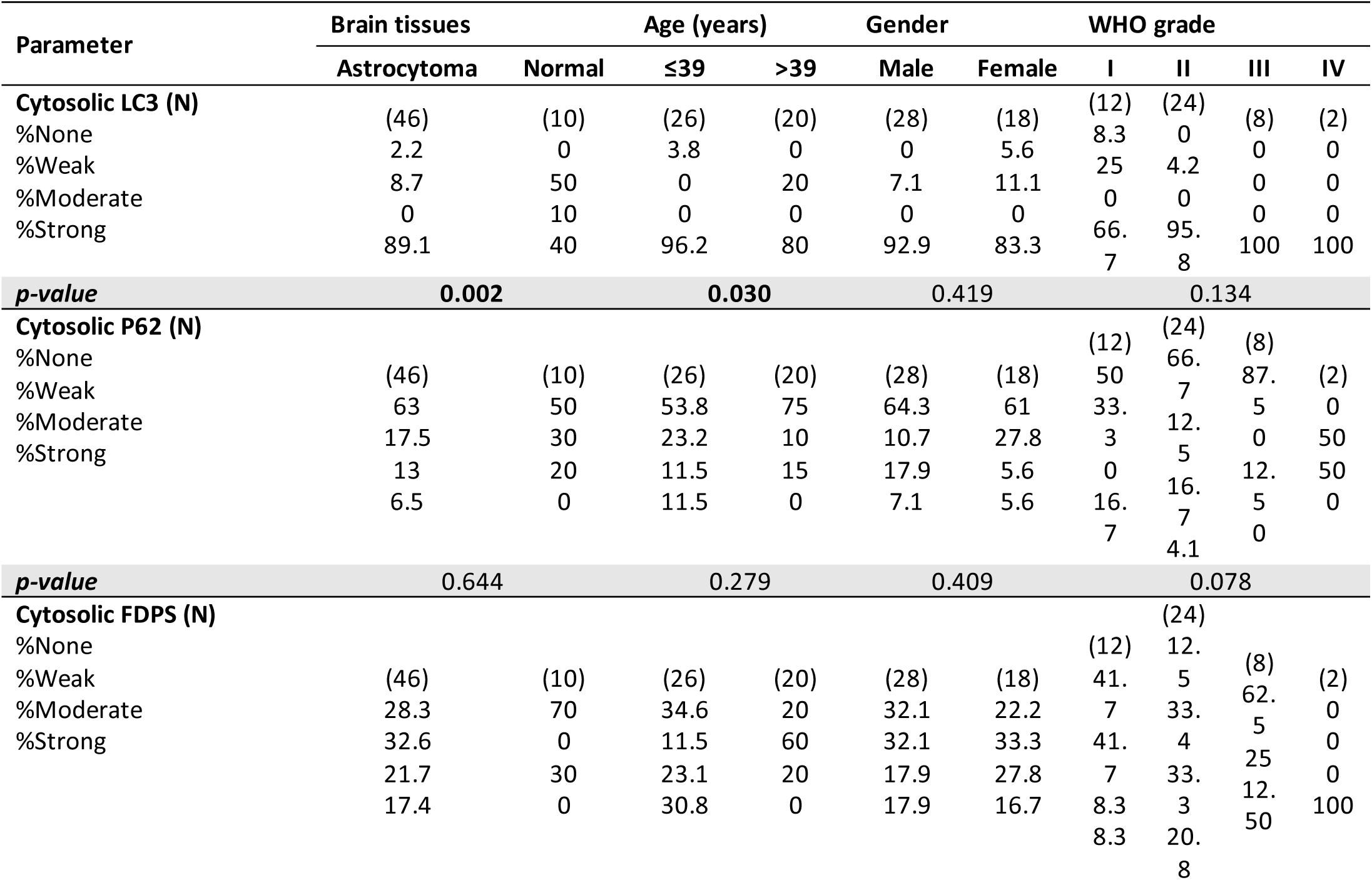
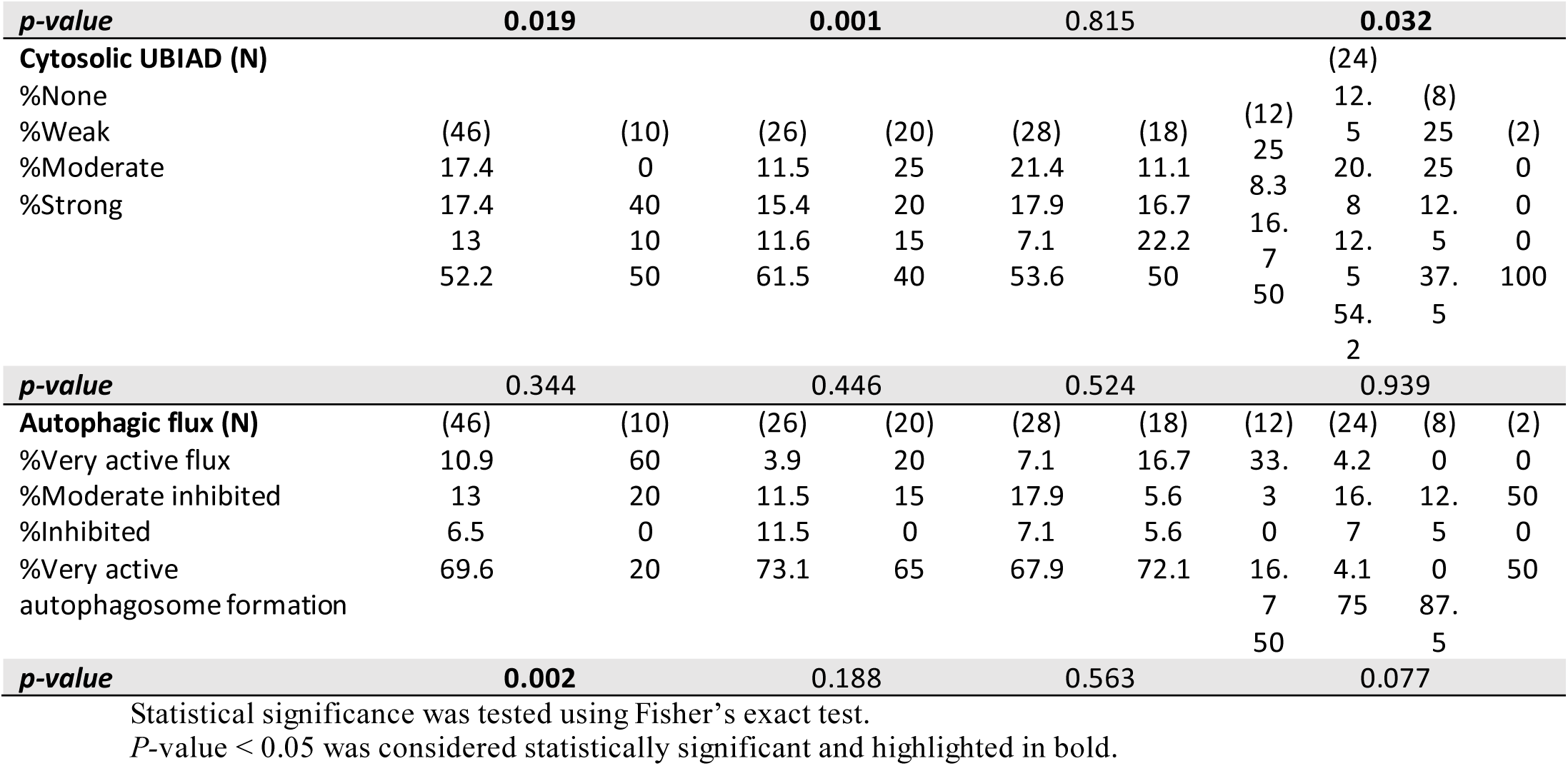
Correlation of cytosolic LC3, P62, FDPS, and UBIAD protein expression and autophagic flux with clinicopathological features of astrocytoma

**Table 3.**
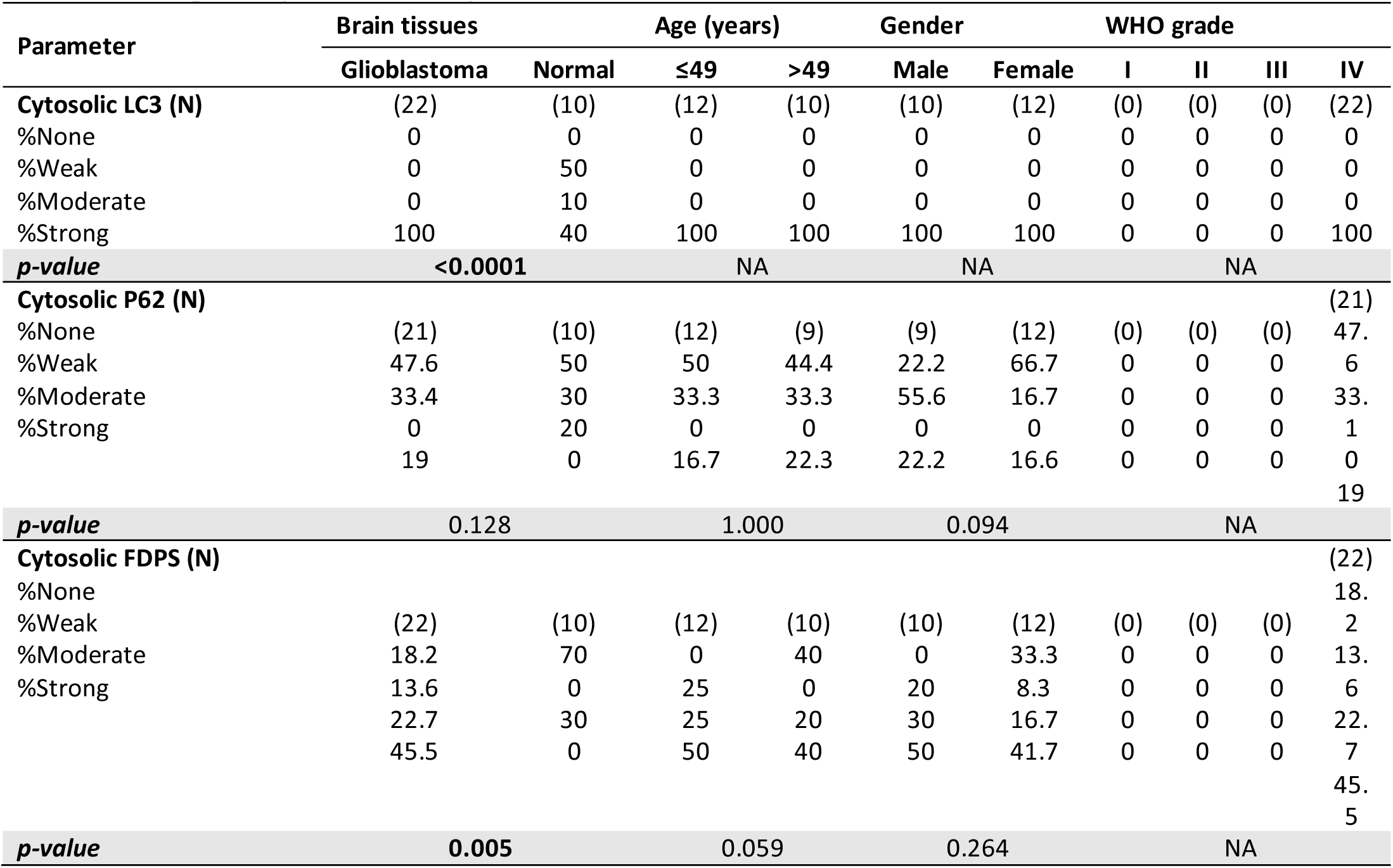
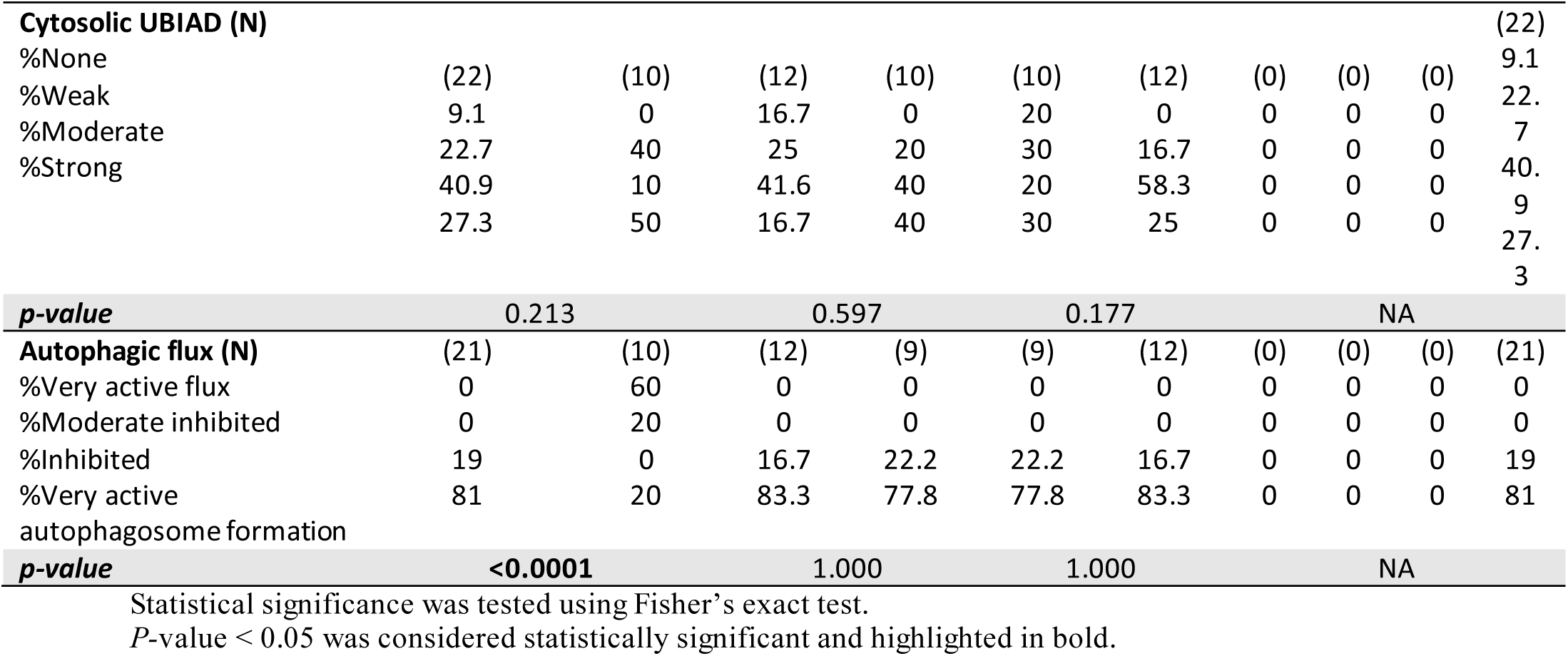
Correlation of cytosolic LC3, P62, FDPS, and UBIAD protein expression and autophagic flux with clinicopathological features of glioblastoma

Correlation analyses revealed coordinated relationships among autophagy-related markers, cholesterol metabolism enzymes, and clinicopathological variables. Spearman correlation matrices demonstrated strong positive associations between LC3β puncta and inferred autophagy activity across all tumors and within astrocytoma and glioblastoma subgroups (**Figure 1E**, **Table 1**). p62 puncta showed a consistent inverse correlation with autophagic flux activity across all tumor subtypes, including glioblastoma (r = −0.739, P < 0.0001), supporting impaired autophagic clearance. LC3β puncta were positively correlated with UBIAD1 expression in all tumors and astrocytoma, whereas FDPS expression correlated significantly with histological grade (P = 0.008) and autophagosome accumulation (P = 0.027).

Collectively, these data demonstrate a progressive remodeling of autophagy and cholesterol metabolism pathways during glioma progression, characterized by enhanced autophagosome accumulation and FDPS upregulation from astrocytoma to glioblastoma. This coordinated autophagy–metabolism dysregulation defines a shared adaptive program associated with malignant progression and suggests that these pathways may also contribute to therapeutic failure in high-grade gliomas[46–50].

Given that chemoresistance remains a major clinical challenge in glioblastoma and is increasingly linked to cellular stress–adaptive programs and metabolic plasticity, we next investigated whether dysregulated autophagy and cholesterol metabolism are associated with resistance to TMZ. To this end, we established mKate-labeled U251 TMZ-resistant (R) cells and systematically compared their autophagy status and cholesterol metabolic profiles with those of TMZ-sensitive (NR) counterparts. We further examined whether targeting cholesterol metabolism could modulate lipid homeostasis and attenuate TMZ resistance.

Notably, TMZ resistance was accompanied by phenotypic and metabolic reprogramming, including epithelial-to-mesenchymal transition (EMT), mitotic quiescence, and a broader metabolic switch, which we investigate in detail in the subsequent sections.

### TMZ Resistance Is Coupled to EMT, Mitotic Quiescence, and Mitochondrial Remodeling

R cells exhibited significantly reduced sensitivity to TMZ-induced cell death at concentrations ≥10 µM, consistent with our previous observations [12]. To determine whether resistance acquisition was accompanied by phenotypic reprogramming, we examined cellular morphology by bright-field microscopy. Compared with NR cells, R cells displayed an elongated, spindle-like morphology (**Figure 2A**) and a marked increase in cytoplasmic vesicular structures (**Figure 2B**), features consistent with epithelial-to-mesenchymal transition (EMT). Immunoblotting confirmed increased expression of the mesenchymal markers N-cadherin and vimentin in R cells (**Figure 2C**).

**Figure 2.**
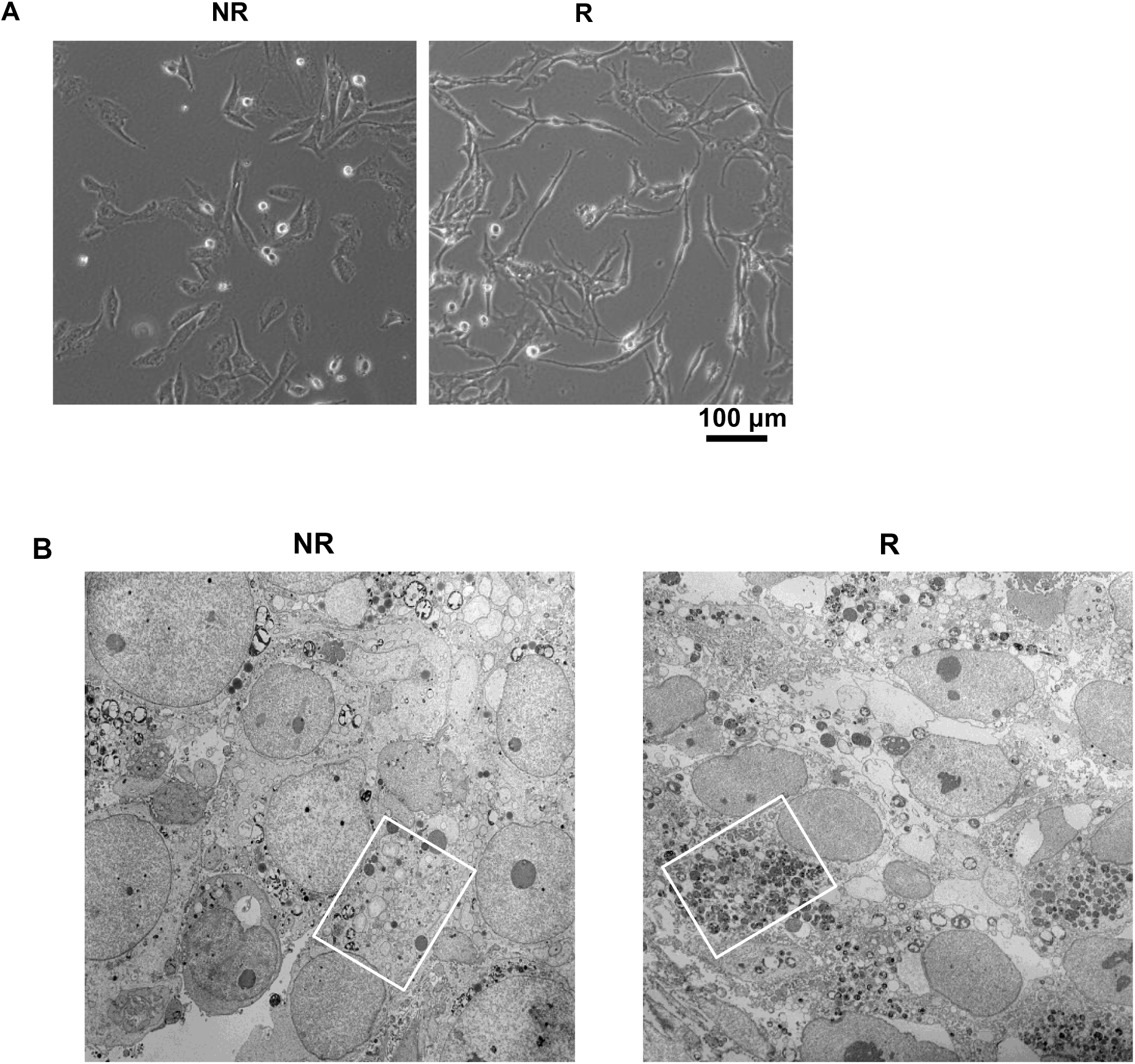
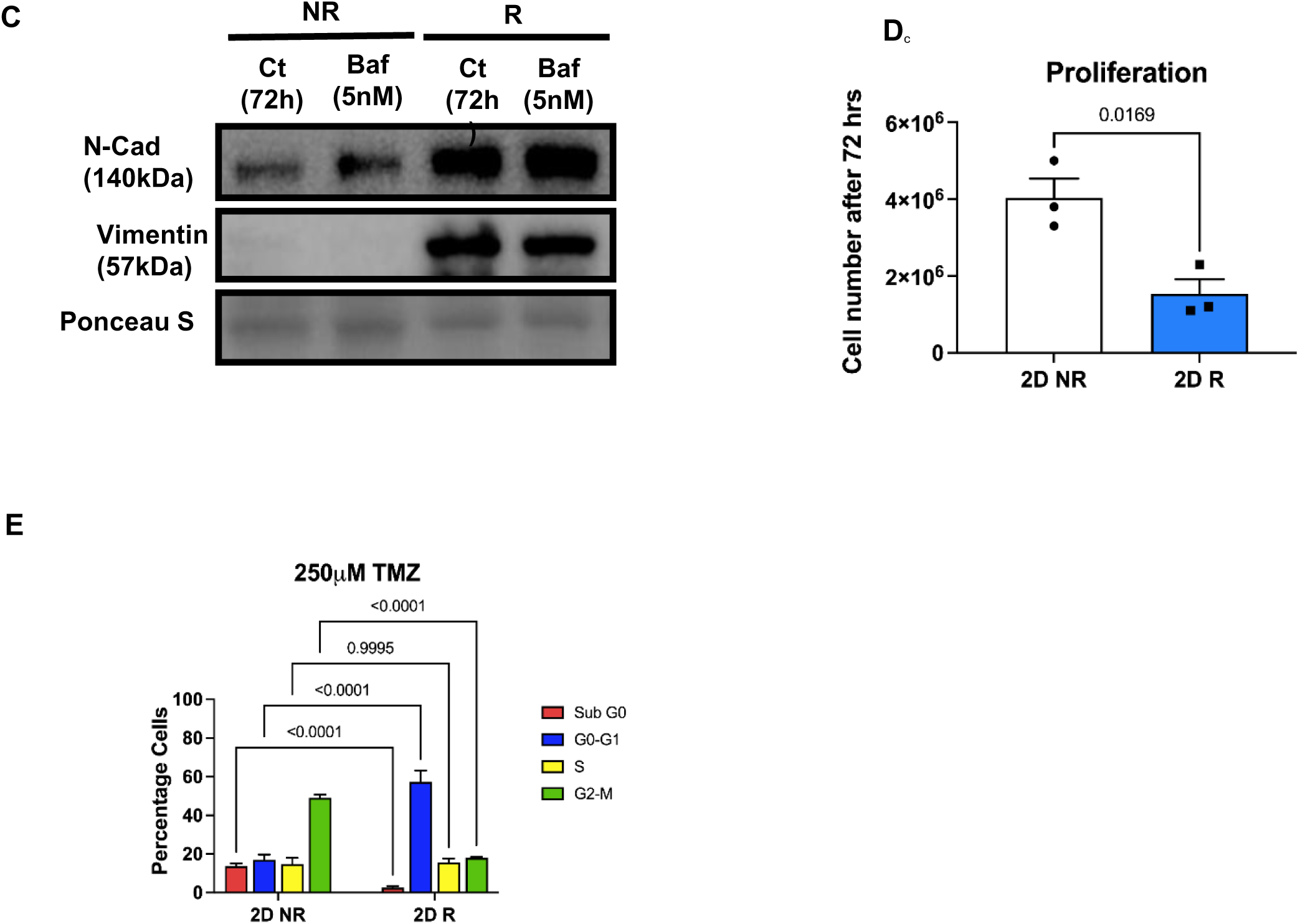
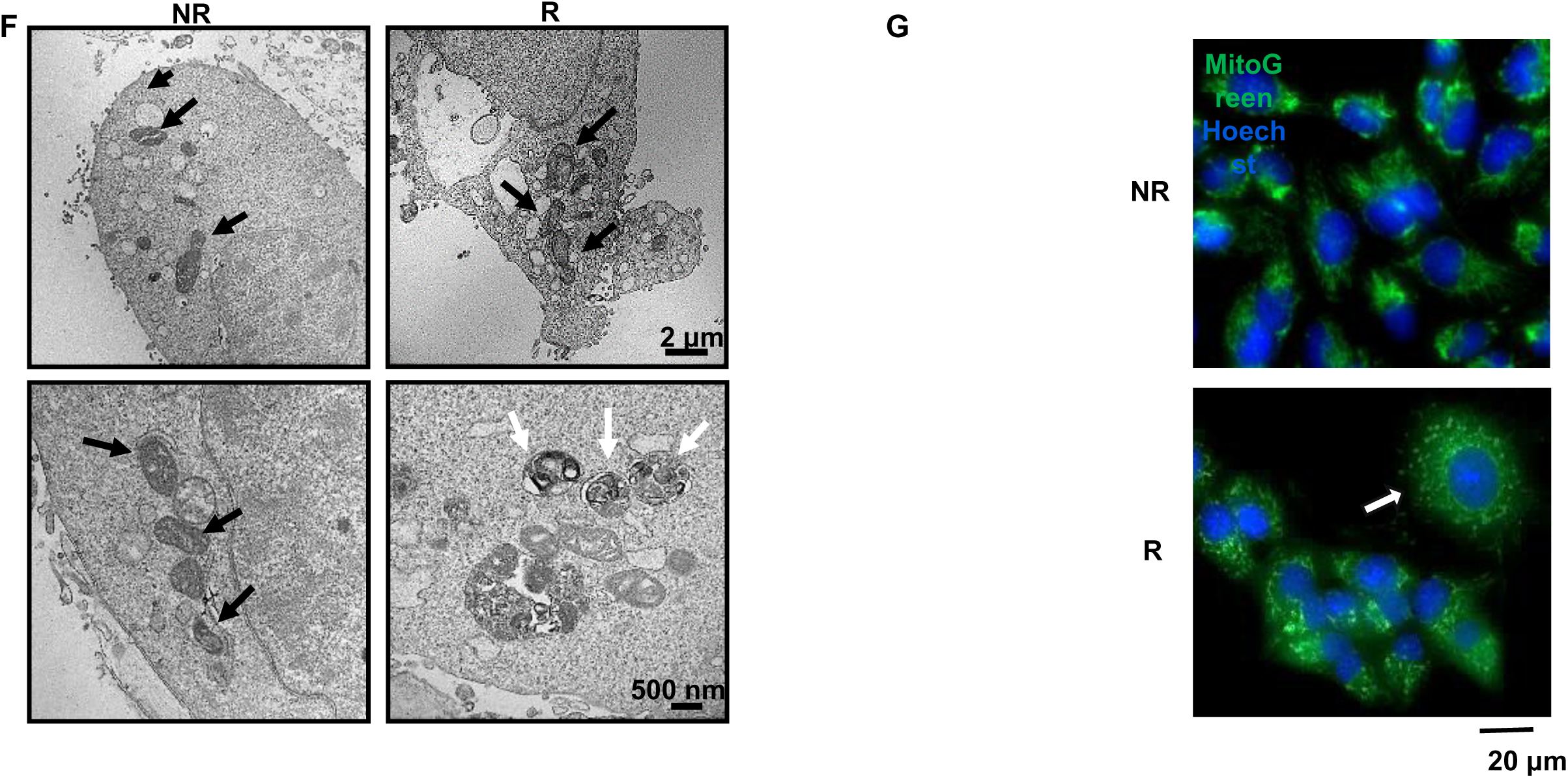
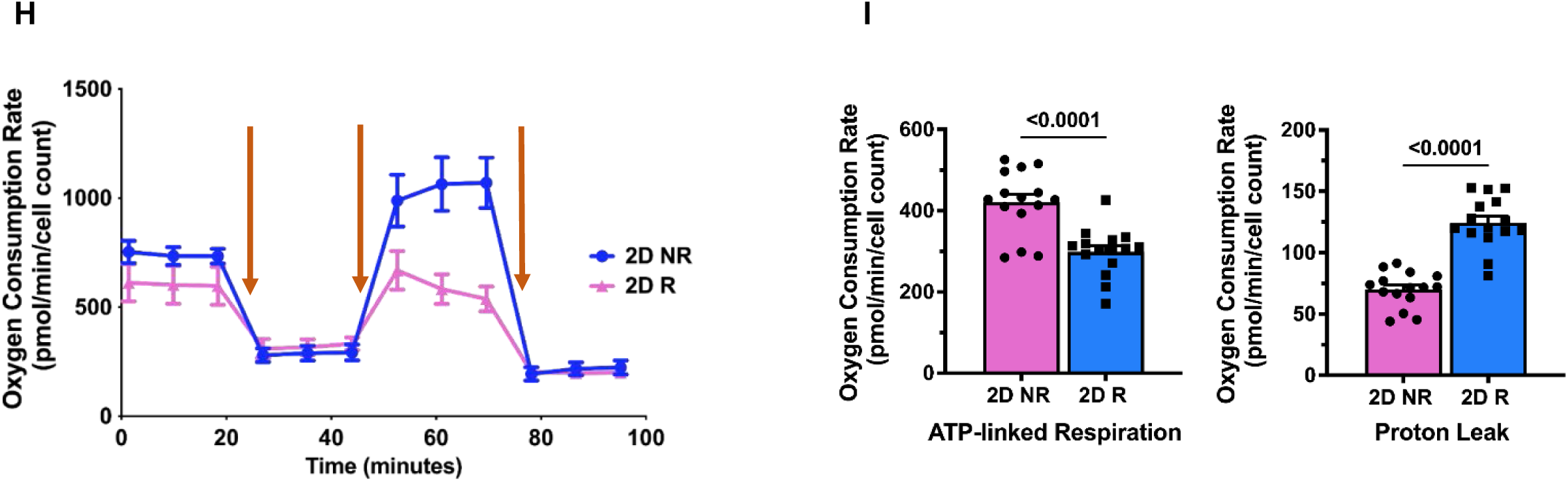
**Stem-like signature of acquired resistance**. (**A**) Bright field imaging comparing R with NR. (**B**) Western blotting of N-cadherin and Vimentin compared between NR and R without treatment and after treatment with Bafilomycin a1 5nM for 72hrs. ( **C**) Proliferation assay comparing the number of the cells 72hrs after seeding 500 cell/ml. (**D**) Flow cytometer results of cell cycle assay comparing R with NR after 48hrs treatment with 250µM TMZ. ( **E**) TEM images compared mitochondrial structure between NR with R. Black arrows show the mitochondria and white arrows show the mitophagosomes. (**F**) Live cell IF imaging of mitochondria using mitoviewGreen, arrow shows mitochondria. (**G**) Oxygen consumption rates (OCR) were analyzed using a Seahorse XFe24 Analyzer and normalized to cell count. Mean ATP production, and proton leak normalized to cell count were calculated. Scale bar=20 um. TEM: The transmission electron microscope. N=3 and 8 biological independent experiment in (**A-F**) and (**G-H**) respectively. ** means *P* ≤ 0.01, *** means *P* ≤ 0.001.

Given the reported links between autophagy and EMT, we next assessed whether acute inhibition of autophagic flux influences mesenchymal marker expression. Treatment with bafilomycin A1 (5 nM, 72 h) did not alter N-cadherin or vimentin levels in either NR or R cells (**Figure 2C**), indicating that maintenance of the mesenchymal phenotype in R cells is not dependent on short-term autophagy flux inhibition.

We then evaluated proliferative dynamics and cell-cycle status. R cells exhibited a substantially reduced proliferation rate, with total cell numbers decreasing to less than half of NR cells after 72 h **(Figure 2D**). Cell-cycle analysis revealed a greater than two-fold reduction in mitotic cells and a three-fold increase in G1-phase arrest in R cells, accompanied by a marked reduction in sub-G0/G1 apoptotic populations (**Figure 2E**), consistent with mitotic quiescence.

Because therapy resistance is frequently associated with mitochondrial adaptation, we examined mitochondrial structure and function. Transmission electron microscopy revealed pronounced mitochondrial abnormalities in R cells, including disrupted cristae architecture and increased mitophagic structures (**Figure 2F**). Live-cell mitochondrial imaging using MitoView Green showed elongated mitochondrial networks in NR cells, whereas R cells exhibited fragmented and clustered mitochondria, consistent with enhanced mitochondrial remodeling ( **Figure 2G**).

To further define metabolic adaptations associated with temozolomide resistance, we assessed mitochondrial respiration in NR and R U251 glioblastoma cells using Seahorse extracellular flux analysis (**Figure 2H-I**). Basal oxygen consumption rates were comparable between NR and R cells; however, following sequential metabolic perturbations, R cells displayed a markedly blunted respiratory response compared with NR cells.

Specifically, NR cells demonstrated a robust increase in respiration following mitochondrial uncoupling, reflecting substantial spare respiratory capacity. In contrast, R cells showed a significantly attenuated response, indicating limited metabolic flexibility under stress conditions (**Figure 2H**). Quantitative analysis confirmed a significant reduction in maximal respiratory capacity and ATP-linked respiration in R cells relative to NR cells, consistent with an altered energetic state (**Figure I**).

Notably, R cells exhibited increased proton leak compared with NR cells (Figure I), suggesting mitochondrial inefficiency and adaptive remodeling of energy utilization rather than enhanced energy production. Together, these findings indicate that acquisition of TMZ resistance is accompanied by a metabolic switch characterized by constrained respiratory reserve and altered mitochondrial stress handling, consistent with a low-proliferative, therapy-tolerant phenotype.

Collectively, these data show that acquisition of TMZ resistance is associated with coordinated phenotypic and bioenergetic reprogramming, characterized by EMT, mitotic quiescence, and mitochondrial remodeling, supporting the existence of a broader metabolic switch that facilitates survival under chemotherapeutic stress.

### Cholesterol Biosynthesis Is Suppressed in TMZ-Resistant Cells in the Context of Impaired Autophagy Flux

Autophagy plays a central role in cellular lipid homeostasis by regulating membrane turnover, lipid droplet utilization, and cholesterol trafficking through autophagosome–lysosome degradation [51]. Impairment of autophagic flux is therefore expected to disrupt cholesterol availability and biosynthetic feedback regulation. Given our previous demonstration that autophagy flux is inhibited in R cells [12], together with the accumulation of autophagosomes and vesicular structures observed in resistant cells, we hypothesized that TMZ resistance is associated with altered cholesterol metabolism as a consequence of defective autophagy-mediated lipid recycling.

Consistent with this hypothesis and our previous findings, ultrastructural analysis by transmission electron microscopy revealed a pronounced increase in intracellular vesicular compartments in R cells, including a higher number of double-membrane vesicles characteristic of autophagosomes, compared with NR cells (**Figure 3A,B**). These findings extend our earlier observations of impaired autophagic clearance and support the presence of disrupted membrane and lipid turnover in resistant cells.

**Figure 3.**
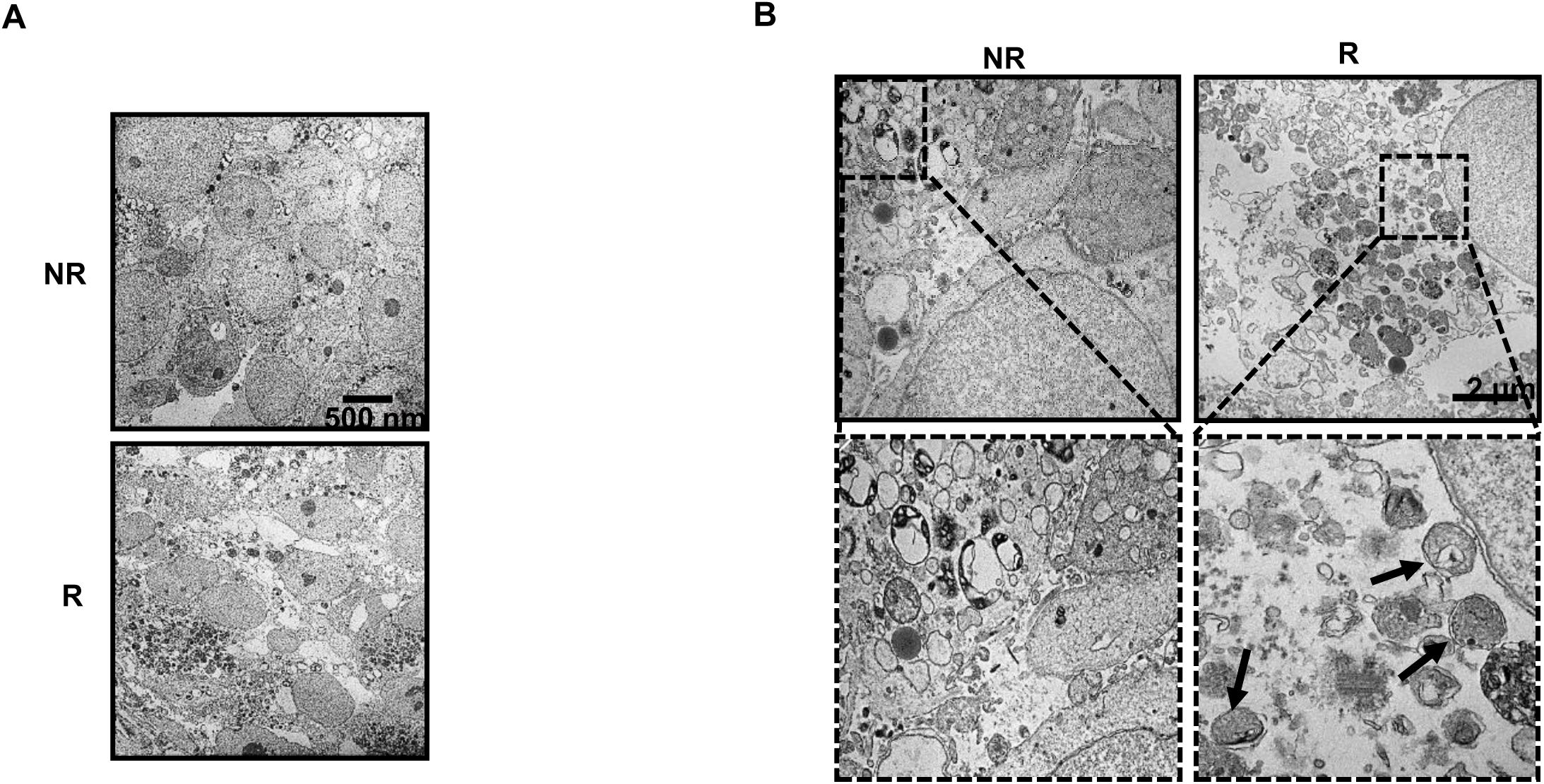
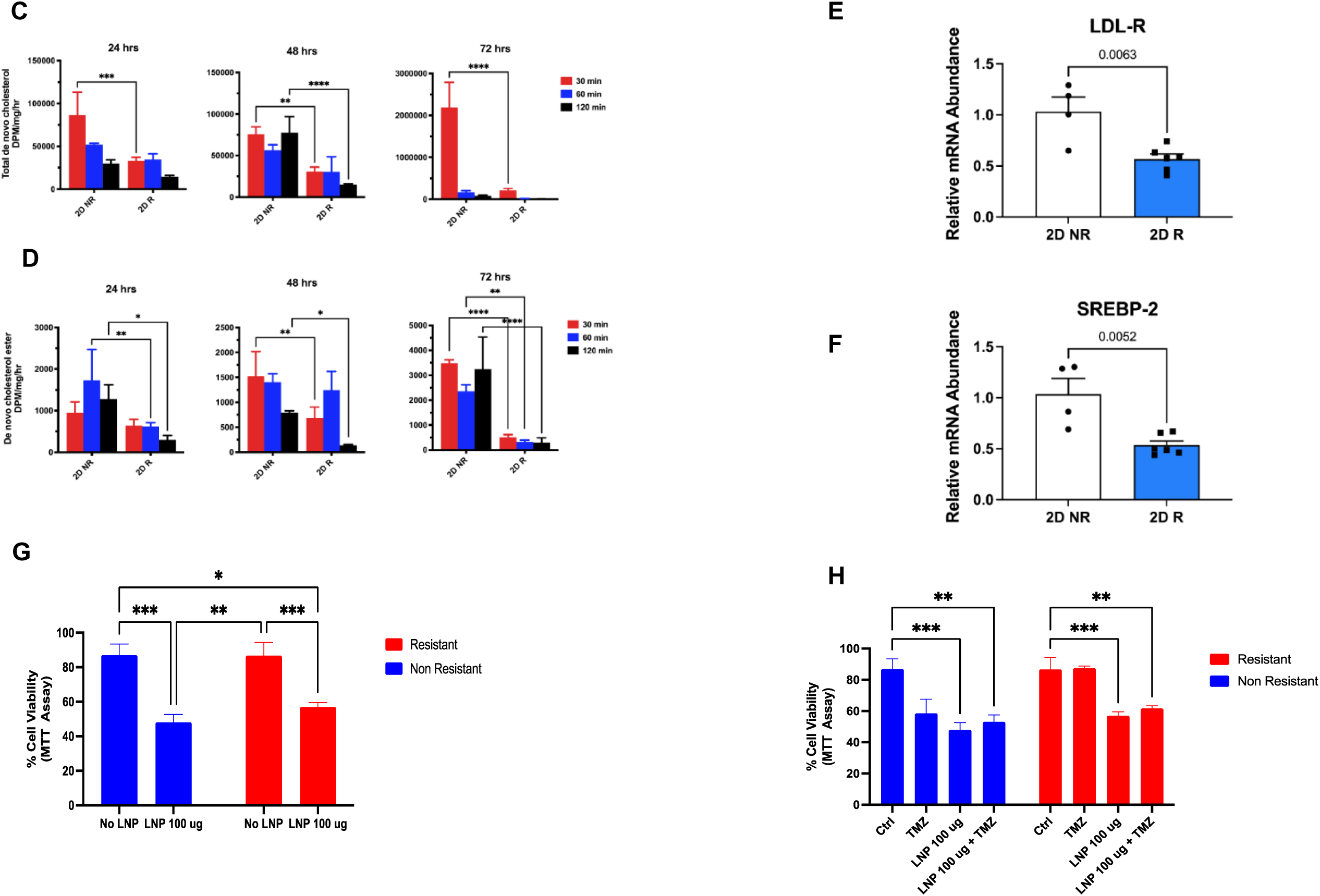
**Autophagy Flux Inhibition and Altered Cholesterol Biosynthesis in Acquired Resistance**. (**A & B**) TEM images compared cellular and intracellular structure between NR with R, arrows show the autophagosomes. (**C**) Total de novo synthesized cholesterol after 24, 48 and 72 hrs culture. It revealed significant decrease in total de novo synthesized cholesterol in R cells in all timepoints. (**D**) De novo Cholesterol esterification after 24, 48 and 72 hrs culture. (**E**) Real-time PCR measured the expression level of LDL-R and (**F**) SREBP-2 genes. LNP induced significant lipooxicity in both NR and R cells (**G**) while did not increase significant change in TMZ-induced cytotoxic effect in R cells (**H**). ** means *P* ≤ 0.01, *** means *P* ≤ 0.001.

Recent multi-omics analyses comparing primary and recurrent glioblastoma specimens have independently identified cholesterol biosynthesis as a pathway significantly altered in TMZ-resistant tumors [52]. Guided by this evidence and our autophagy findings, we quantified total de novo cholesterol synthesis and cholesterol esterification in R and NR cells. Both processes were significantly reduced in R cells across all examined time points ( **Figure 3C,D**), indicating suppression of cholesterol biosynthetic capacity in the resistant state.

To determine whether reduced cholesterol synthesis reflected transcriptional reprogramming, we assessed the expression of LDL receptor (LDL-R), which mediates cholesterol uptake, and sterol regulatory element–binding protein 2 (SREBP2), a master regulator of cholesterol biosynthesis. Real-time PCR analysis revealed significantly decreased LDL-R and SREBP2 mRNA levels in R cells compared with NR cells (**Figure 3E,F**), consistent with impaired cholesterol homeostasis and reduced sterol-driven feedback signaling.

Because impaired autophagy flux may limit endogenous cholesterol recycling, we next tested whether exogenous cholesterol delivery could influence cell viability or TMZ sensitivity. Treatment with lipid nanoparticles (LNPs), which contain approximately 45% cholesterol by mass, induced lipotoxicity in both NR and R cells (**Figure 3G**). However, LNP treatment did not significantly enhance TMZ-induced cytotoxicity in R cells (**Figure 3H**), indicating that cholesterol supplementation alone is insufficient to reverse established TMZ resistance.

Collectively, these data demonstrate that impaired autophagy flux in TMZ-resistant cells is accompanied by suppression of cholesterol biosynthesis and esterification, together with transcriptional downregulation of cholesterol regulatory pathways. This coordinated dysregulation links defective autophagy-mediated lipid recycling to altered cholesterol metabolism and supports a model in which metabolic adaptation contributes to the maintenance of TMZ resistance.

### Simvastatin Fails to Resensitize TMZ-Resistant Cells to TMZ-Induced Apoptosis

Our previous work demonstrated that simvastatin (Simva) sensitizes glioblastoma cells to temozolomide (TMZ)-induced apoptosis through inhibition of autophagic flux [36]. Given our current findings that TMZ-resistant (R) cells exhibit impaired autophagy flux, suppressed cholesterol biosynthesis, and extensive metabolic reprogramming, we next asked whether Simva could overcome TMZ resistance in this context.

R cells were treated with Simva alone or in combination with TMZ, and apoptotic responses were assessed. In contrast to TMZ-non-resistant (NR) cells, neither Simva monotherapy nor combined Simva–TMZ treatment induced a significant increase in apoptotic cell death in R cells ( **Figure 4A**). Consistently, caspase-3/7 activity (**Figure 4B**) and caspase-9 activation (**Figure 4C**) were not significantly elevated in R cells following either treatment compared with NR cells.

**Figure 4.**
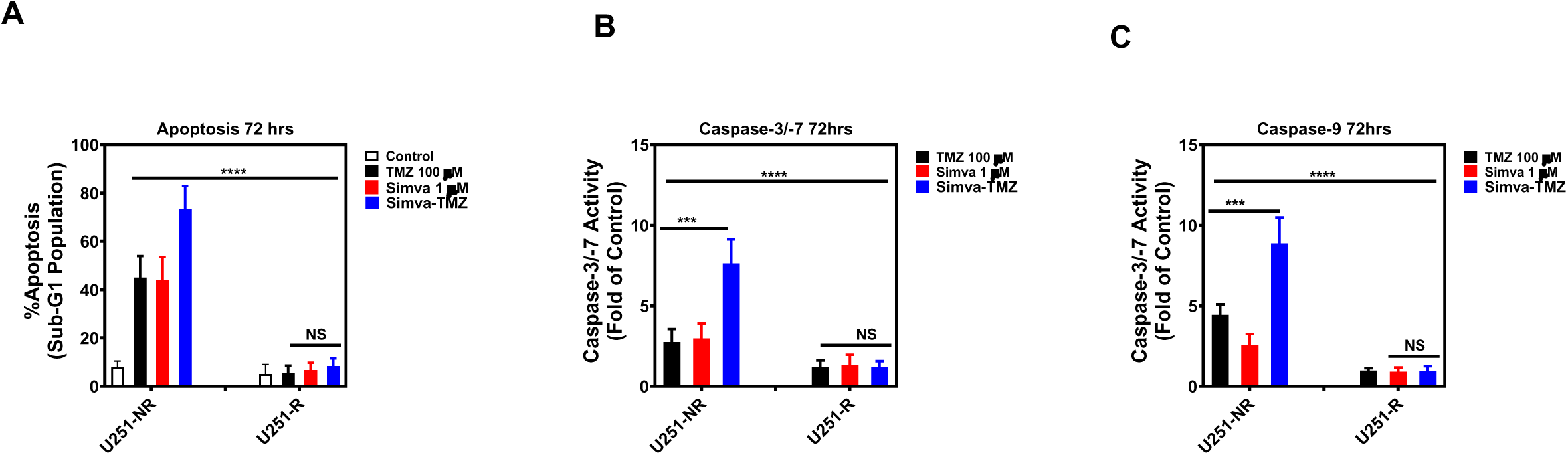
**Simva and TMZ-Simva treatment do not induces apoptosis in U251 R cells**. U251 NR and R cells were treated with TMZ (100 µM), Simva (1 µ) and TMZ-Simva for 72 h. Apoptosis was evaluated using propidium idodide Nicoletti assay and caspase-3/-7 and -9 luminometric assay. TMZ, Simva and TMZ-Simva treatment did not induce any apoptosis (**A**), caspase-3/-7 (**B**), and caspase-9 activation (**C**). All experiments were done in 3 independent biological replicates. ** means *P* ≤ 0.01, *** means *P* ≤ 0.001.

These results indicate that, unlike in TMZ-sensitive cells, inhibition of the mevalonate pathway by Simva is insufficient to restore TMZ-induced apoptotic signaling in the resistant state. Together with our earlier data, these findings suggest that the autophagy–cholesterol axis in R cells is rewired in a manner that uncouples statin-mediated autophagy modulation from apoptotic execution, highlighting the robustness of the resistant phenotype.

### Simvastatin Exposure Reveals Cholesterol Metabolic Plasticity in TMZ-Resistant Glioblastoma

Our preceding analyses demonstrated that R glioblastoma cells exhibit impaired autophagy flux, accumulation of autophagosomes, and suppression of endogenous cholesterol biosynthesis, together with resistance to statin-mediated apoptotic sensitization. Because cholesterol homeostasis is tightly regulated by both de novo synthesis and autophagy-dependent lipid recycling, these findings raised the possibility that pharmacologic inhibition of the mevalonate pathway may differentially impact cholesterol handling in resistant versus non-resistant cells.

Simvastatin (ST), a clinically approved HMG-CoA reductase inhibitor, directly targets the rate-limiting step of cholesterol biosynthesis and has been shown to modulate autophagy and membrane lipid composition in cancer cells [53, 54]. On the other hand, Acquired resistance in glioblastoma is a significant hurdle in effective treatment, often associated with metabolic reprogramming, including changes in cholesterol metabolism [12, 55, 56]. We therefore used ST as a metabolic probe to interrogate how TMZ resistance alters cholesterol metabolic flexibility and compensatory lipid remodeling. By comparing the effects of ST, TMZ, and combined TMZ–ST treatment on cholesterol profiles in NR and R U251 cells, we aimed to define resistance-specific adaptations in cholesterol metabolism that persist despite pharmacologic pathway inhibition. To this end, we performed comprehensive lipidomic analyses of free cholesterol and cholesterol esters to assess how R and NR cells differentially remodel cholesterol pools in response to ST-based perturbation.

A comprehensive heatmap analysis revealed significant alterations in cholesterol profiles, including free cholesterol and a range of cholesterol esters (CE 14:0 to CE 22:6), between NR and R glioblastoma cells under various treatment conditions (**Figure 5A**). Principal component analysis (PCA) of cholesterol species demonstrated consistent and robust segregation between **TMZ-resistant (R)** and **NR** groups across all treatment conditions (B–E). In panel **B**, R samples were clearly separated from NR samples primarily along PC1, which explained **73.9%** of the total variance, with PC2 accounting for **17.5%**, indicating marked baseline differences in cholesterol-associated profiles. Under **ST treatment** (**C**), this separation was maintained, with **PC1 explaining 77.1%** and **PC2 15.8%** of the variance, and R-ST samples forming a compact cluster distinct from NR-ST samples. Under **TMZ treatment** (**D**), PCA revealed pronounced segregation between R-TMZ and NR-TMZ groups, with **PC1 and PC2 accounting for 73.4% and 19.6%** of the variance, respectively, and R-TMZ samples clustering tightly. In the **ST/TMZ combination treatment** (**E**), R-ST/TMZ samples remained clearly separated from NR-ST/TMZ samples, demonstrating that TMZ resistance is associated with a distinct cholesterol species signature that persists and is further accentuated under combination therapy.Volcano plot analysis revealed that, in control groups, CE 22:0 levels were significantly increased in R cells compared to NR cells, indicating inherent differences in cholesterol metabolism associated with resistance (**Figure 5F**). Under ST treatment, several cholesterol esters, including CE 22:6, CE 22:4, and CE 22:5, were significantly upregulated in R cells, with amplified changes observed in R cells, indicating a resistance-mediated adaptation to ST treatment (**Figure 5G**). Interestingly, TMZ treatment alone did not result in significant changes in cholesterol esters, suggesting limited effects of TMZ on cholesterol metabolism (**Figure 5H**). However, combined TMZ-ST treatment led to a significant upregulation in multiple cholesterol esters, including CE 22:5, CE 22:4, and CE 22:6, with the most pronounced effects observed in R cells, highlighting the synergistic potential of this combination (**Figure 5I**). Quantitative analyses of cholesterol components, including free cholesterol and specific esters (CE 16, CE 17, CE 18, CE 20, and CE 22), confirmed these trends, with R cells consistently exhibiting higher levels compared to NR cells across all treatment groups (except CE 18:1), particularly under combined TMZ-ST therapy (**Figure 5J-P**). Our findings reveal that TMZ resistance in GB cells is associated with significant alterations in cholesterol metabolism. While ST and the ST-TMZ combination disrupted CE profiles, neither treatment induced apoptosis or activated Caspase-3/-7 or Caspase-9 in R cells compared to NR cells. Our KEGG analysis (**Figure 5Q and R**) revealed that the top five pathways associated with differences between R and NR GB cells were Steroid hormone biosynthesis, Ovarian steroidogenesis, Neuroactive ligand-receptor interaction, Insulin resistance, and Metabolic pathways. These pathways are intricately linked to autophagy, providing a strong rationale for its investigation in validating our lipidomics results. Steroid hormone biosynthesis influences autophagy through glucocorticoids, which modulate autophagic flux by interacting with key pathways such as mTOR [57]. Similarly, Ovarian steroidogenesis involves estrogen and progesterone, which regulate autophagy in various cellular contexts, including cancer [58, 59]. The Neuroactive ligand-receptor interaction pathway highlights neurotransmitters and receptors that impact autophagy via calcium signaling and neurotrophic factors [60]. Insulin resistanc**e** directly alters autophagy by disrupting glucose metabolism and activating stress responses critical for cellular adaptation [61]. Lastly, Metabolic pathways broadly integrate lipid and energy metabolism with autophagic processes to regulate lipid turnover and maintain homeostasis [62]. These findings underscore the significant role of autophagy in these pathways, justifying its investigation in the next section as a biological validation of our lipidomics findings.

**Figure 5.**
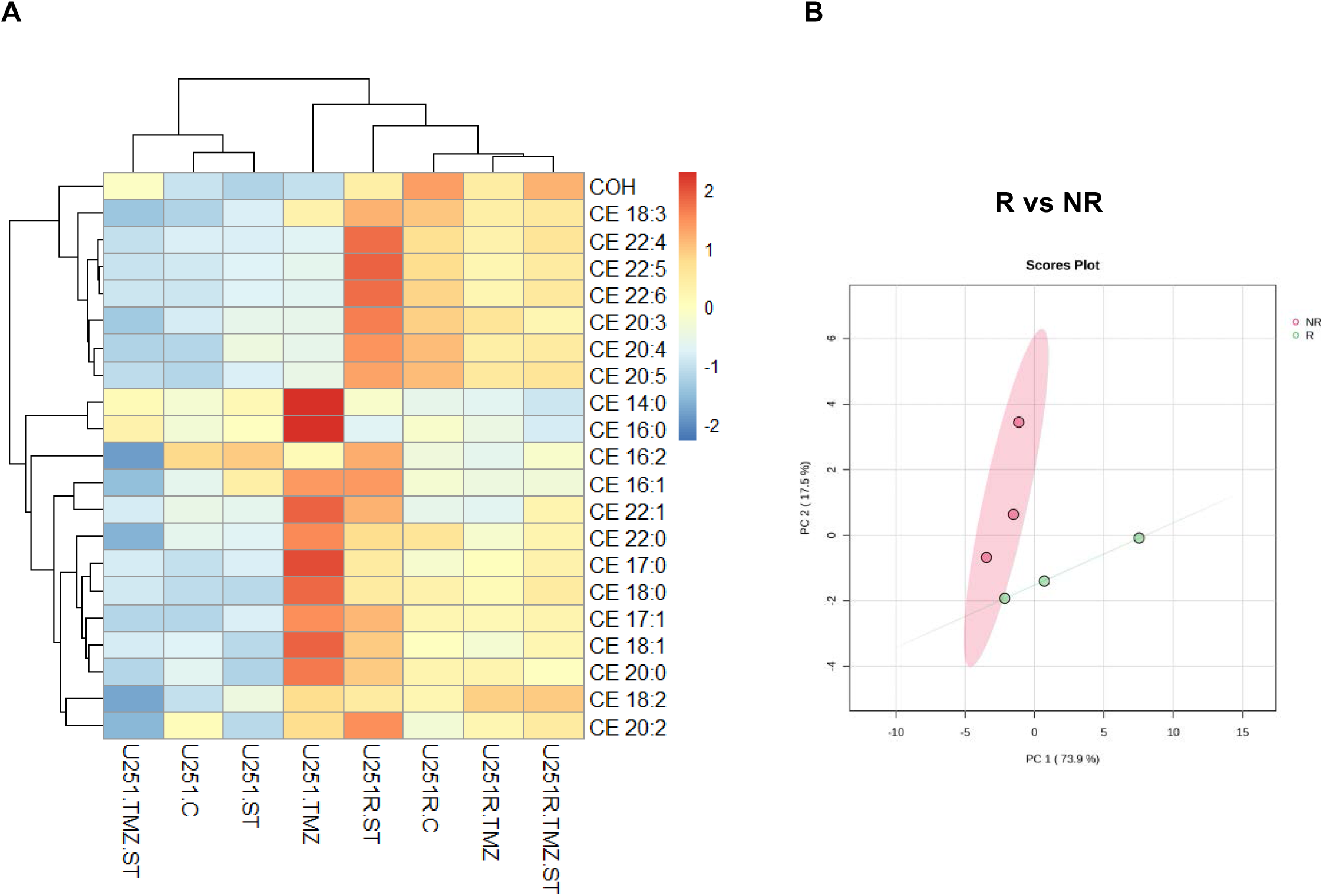
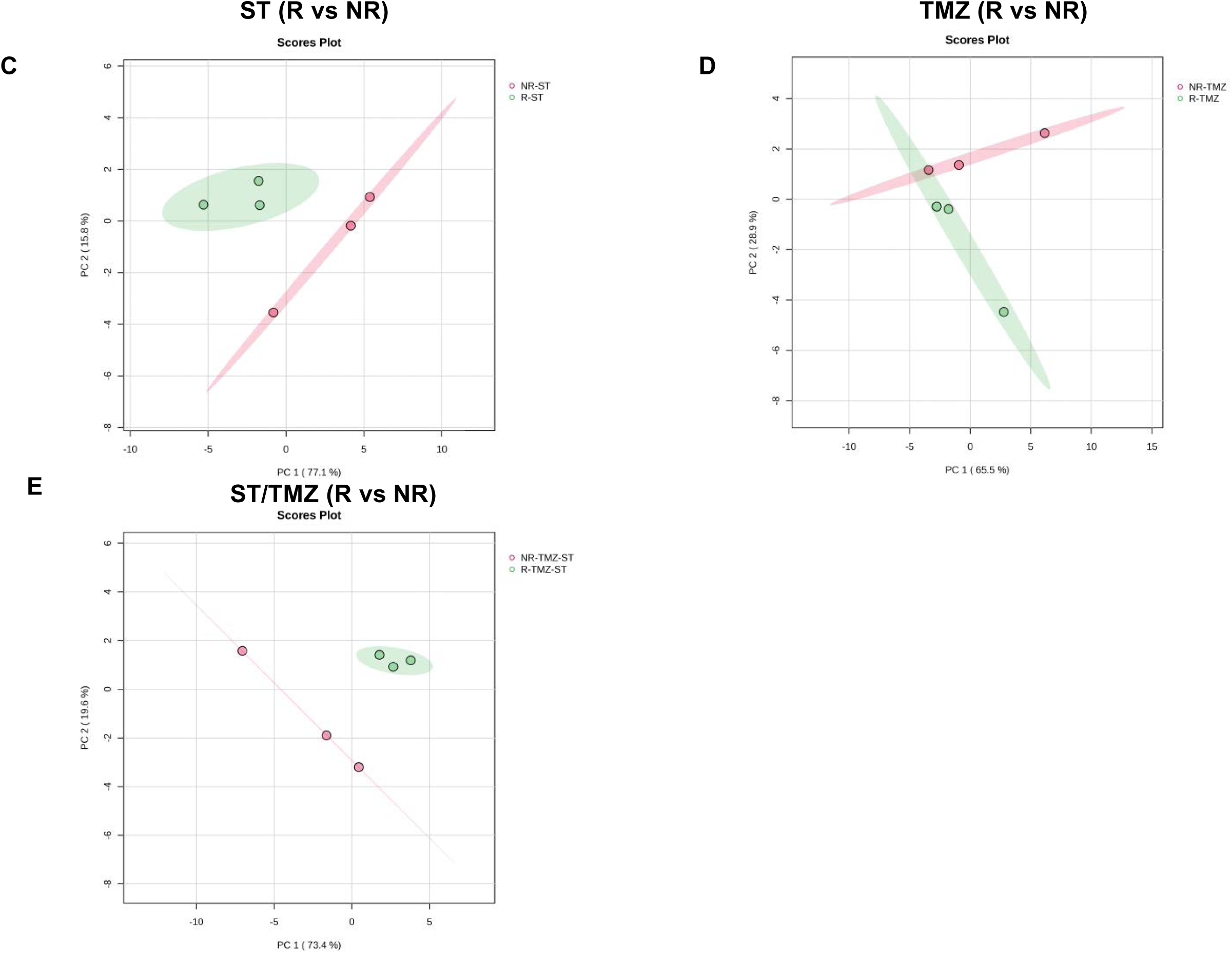
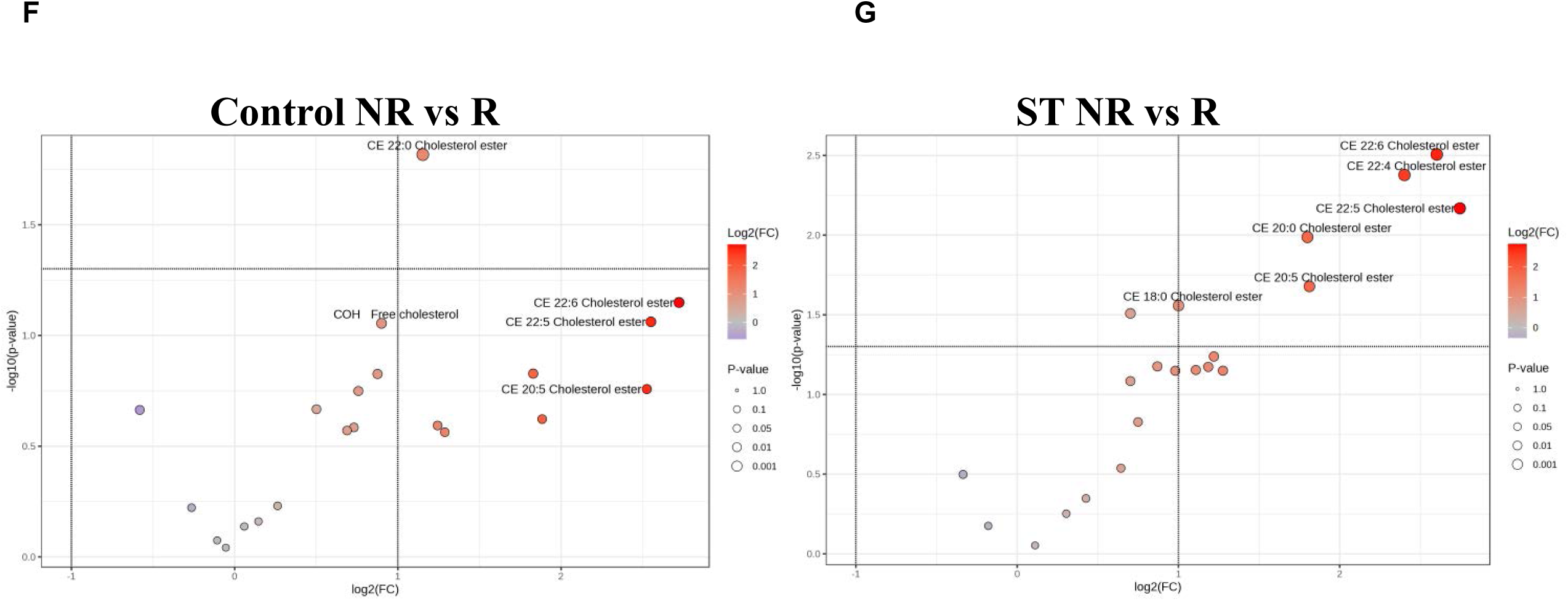
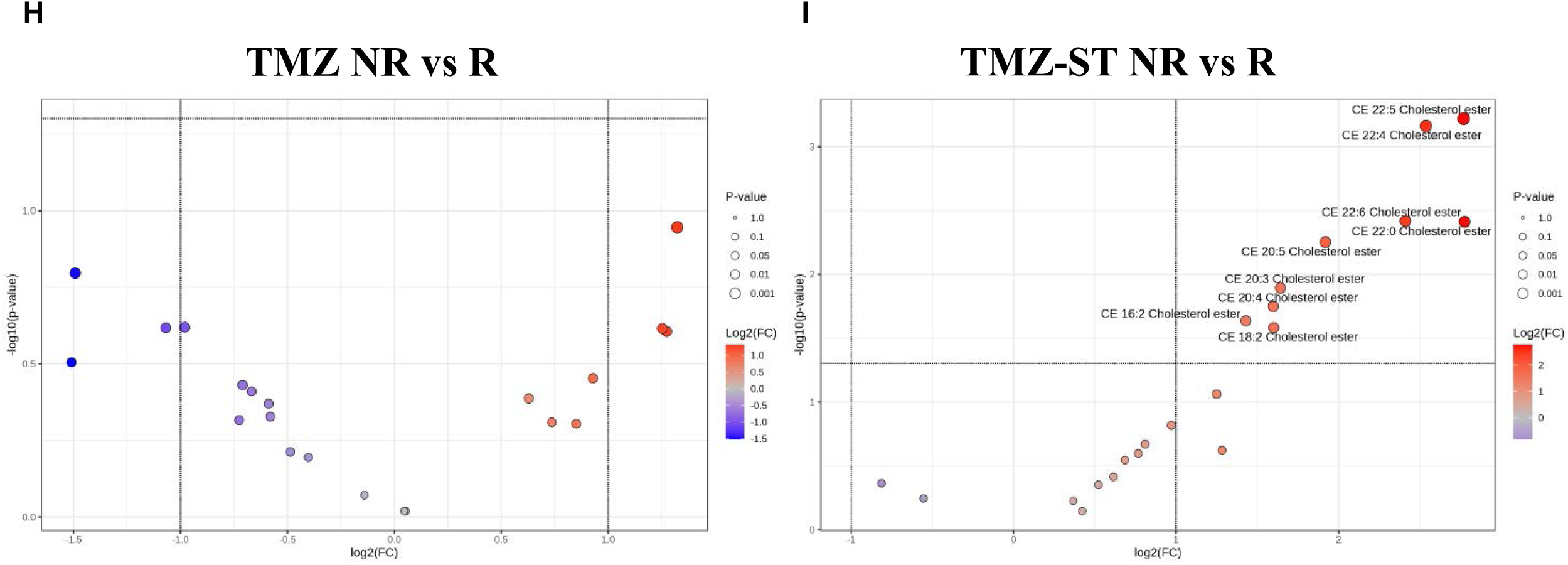
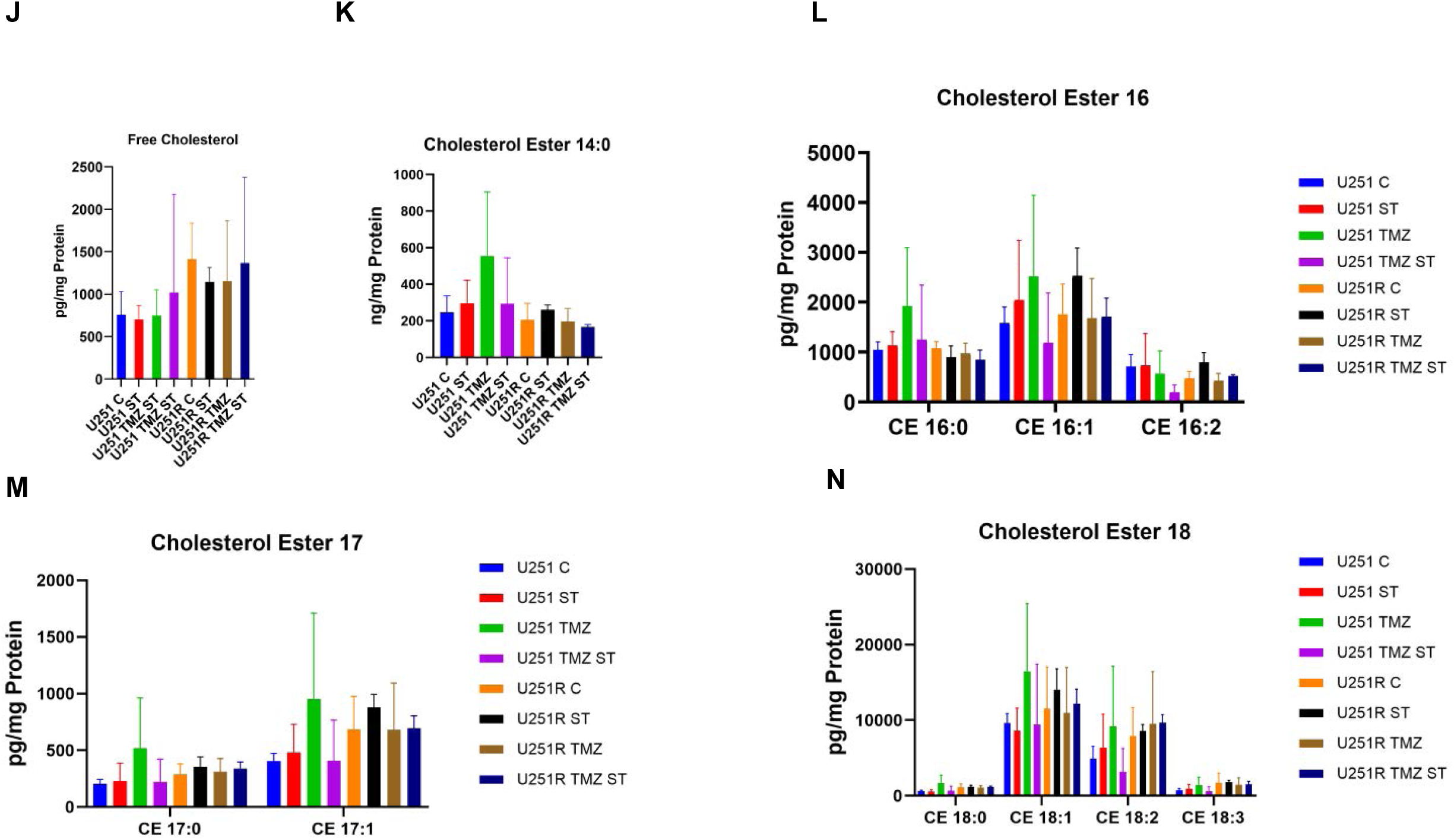
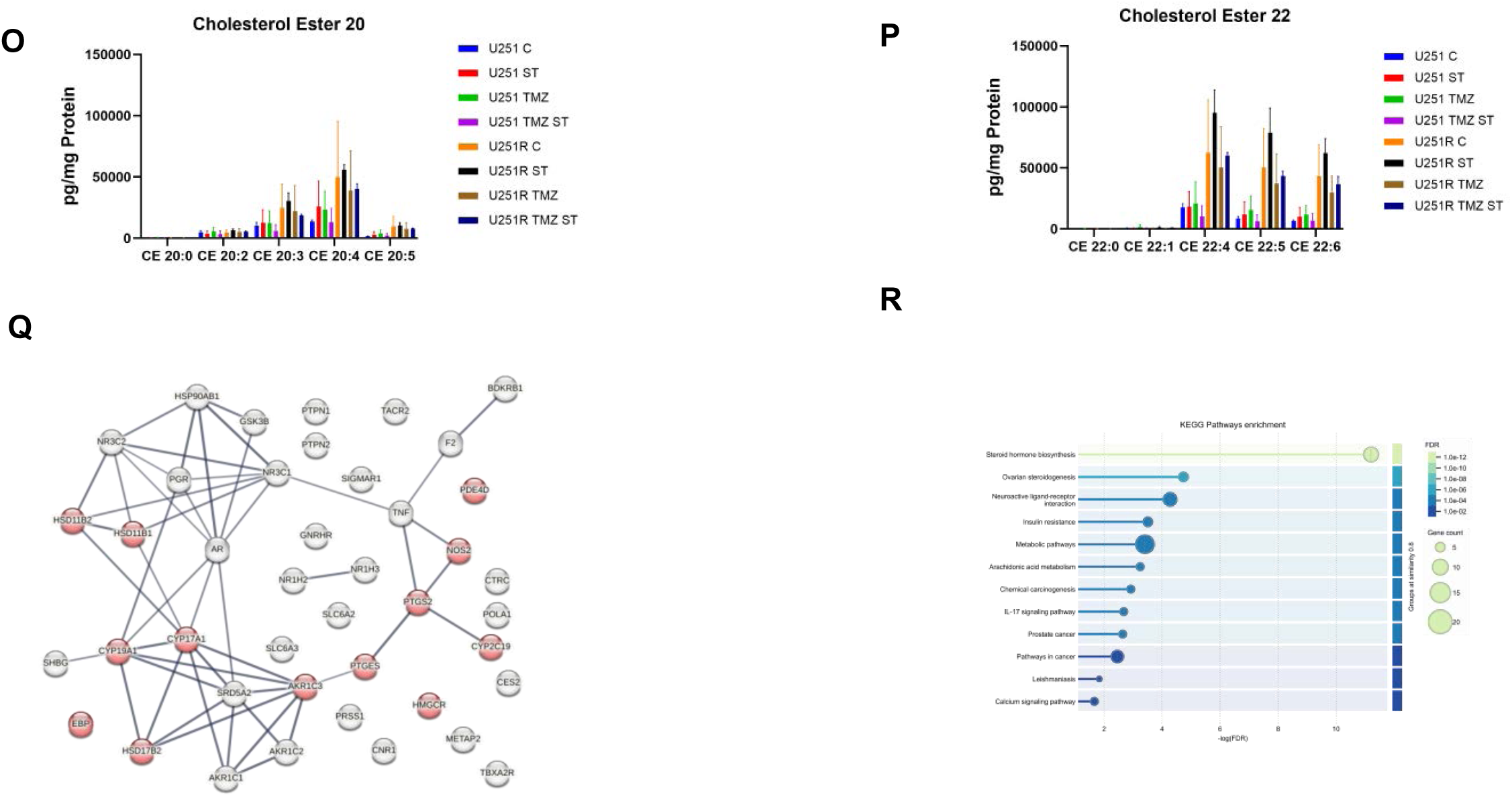
Impact of Temozolomide (TMZ), Simvastatin (ST), and Combined TMZ-ST Treatment on Cholesterol Metabolism in U251 Non-Resistant NR and R Glioblastoma Cells. **A)** Heatmap generated using pheatmap illustrating the effects of Temozolomide (TMZ), Simvastatin (ST), and combined TMZ-ST treatment on cholesterol metabolism in U251 Non-Resistant (NR) and Resistant (R) glioblastoma cells. The heatmap includes free cholesterol and a series of cholesterol esters, ranging from CE 14:0 to CE 22:6, to highlight changes in cholesterol profiles under different treatment conditions (average of n=3). Principal component analysis (PCA) of cholesterol species showing separation between R and NR groups under different treatment conditions. Panels **B–E** represent baseline, ST, TMZ, and ST/TMZ combination treatments, respectively. The percentage of variance explained by PC1 and PC2 is indicated on each axis. Distinct clustering highlights treatment-dependent differences in cholesterol species associated with TMZ resistance. **F-I)** Volcano plots generated through MetaboAnalyst 6.0 illustrating differential expression of cholesterol species (Free Cholesterol and Cholesterol Esters) in U251 R cells compare to U251 NR cells. Comparisons include control vs. treatments with Simvastatin (ST), Temozolomide (TMZ), and combined TMZ-ST. Treatments are designated as follows: control (no treatment) (F), ST (G), TMZ (H), and TMZ-ST (I). Each treatment has done for three replicates (n=3). In the comparison between NR and R control groups, the volcano plot highlights an increase in CE 22:0 cholesterol ester levels in the resistant group, indicating differential cholesterol metabolism associated with treatment resistance in U251 glioblastoma cells (F). In the ST treatment comparing NR vs R cells, the volcano plot highlights several cholesterol esters that are significantly altered. CE 22:6, CE 22:4, and CE 22:5 are the most significantly downregulated upregulated cholesterol esters, suggesting their levels are substantially lower higher in resistant (R) cells compared to non-resistant (NR) cells when treated with Simvastatin. CE 20:0 and CE 20:5 also show notable up regulation downregulation. In contrast, CE 18:0 does not exhibit as strong a change as the others. These alterations indicate a distinct lipidomic profile response to ST treatment between NR and R U251 glioblastoma cells (H). Volcano plot showing the effects of Temozolomide (TMZ) treatment on cholesterol ester profiles in U251 NR and R cells. The analysis reveals no meaningful changes in the levels of cholesterol esters, indicating a lack of differential expression between treated and control groups for the compounds analyzed (n=3) (G). Volcano plot illustrating changes in cholesterol ester profiles between NR and R U251 cells following TMZ-ST treatment. The plot shows an increase in the levels of CE 22:5, CE 22:4, CE 22:6, CE 22:0, CE 20:5, CE 20:3, CE 20:4, CE 16:2, and CE 18:2 in R cells compared to NR cells, suggesting a differential lipid metabolic response to combined drug treatment (n=3) (J). **J-P)** Concentration of cholesterol family components (Free Cholesterol, CE 14, CE 16, CE 17, CE 18, CE 20, CE 22) in U251 NR and R cells treated with ST, TMZ, and TMZ-ST that generated by GraphPad Prism 10. This comparison highlights the differential lipid metabolic responses to the treatments across the two cell lines. **Q,R)** Subpanel Q andR shows a protein–protein interaction (PPI) network constructed from genes associated with cholesterol metabolic remodeling in R cells following TMZ, ST, and combined TMZ–ST treatment. Nodes represent individual proteins and edges denote known or predicted interactions. Proteins highlighted in red indicate highly connected hub nodes within the network, including key enzymes involved in cholesterol and steroid metabolism (e.g., AKR1C family members, HSD17B enzymes, and cytochrome P450–related proteins). The presence of these hubs within a densely interconnected network supports coordinated metabolic and stress-adaptive signaling underlying cholesterol plasticity and drug resistance in TMZ-resistant glioblastoma cells**. S**) Subpanel S depicts pathway enrichment analysis of genes associated with cholesterol metabolism in U251 R cells following TMZ, ST, and combined TMZ–ST treatment. The top enriched pathways include Steroid hormone biosynthesis, Ovarian steroidogenesis, Neuroactive ligand–receptor interaction, Insulin resistance, and Metabolic pathways. Color intensity reflects adjusted P values, while bubble size indicates the number of genes contributing to each pathway. These enrichments support coordinated metabolic and stress-adaptive reprogramming underlying cholesterol plasticity and drug resistance.

### Simvastatin and Temozolomide Fail to Restore Autophagy Flux in TMZ-Resistant Glioblastoma Cells

To functionally validate our lipidomic findings and determine whether pharmacologic perturbation of cholesterol metabolism influences autophagy in the resistant state, we examined the effects of temozolomide (TMZ; 100 µM), simvastatin (Simva; 1 µM), and combined TMZ–Simva treatment on autophagy flux in U251 NR and R glioblastoma cells over 72 hours. Autophagy flux was assessed using transmission electron microscopy (TEM) and immunoblot analysis of LC3β and p62/SQSTM1 (**Figure 6A,B**).

**Figure 6.**
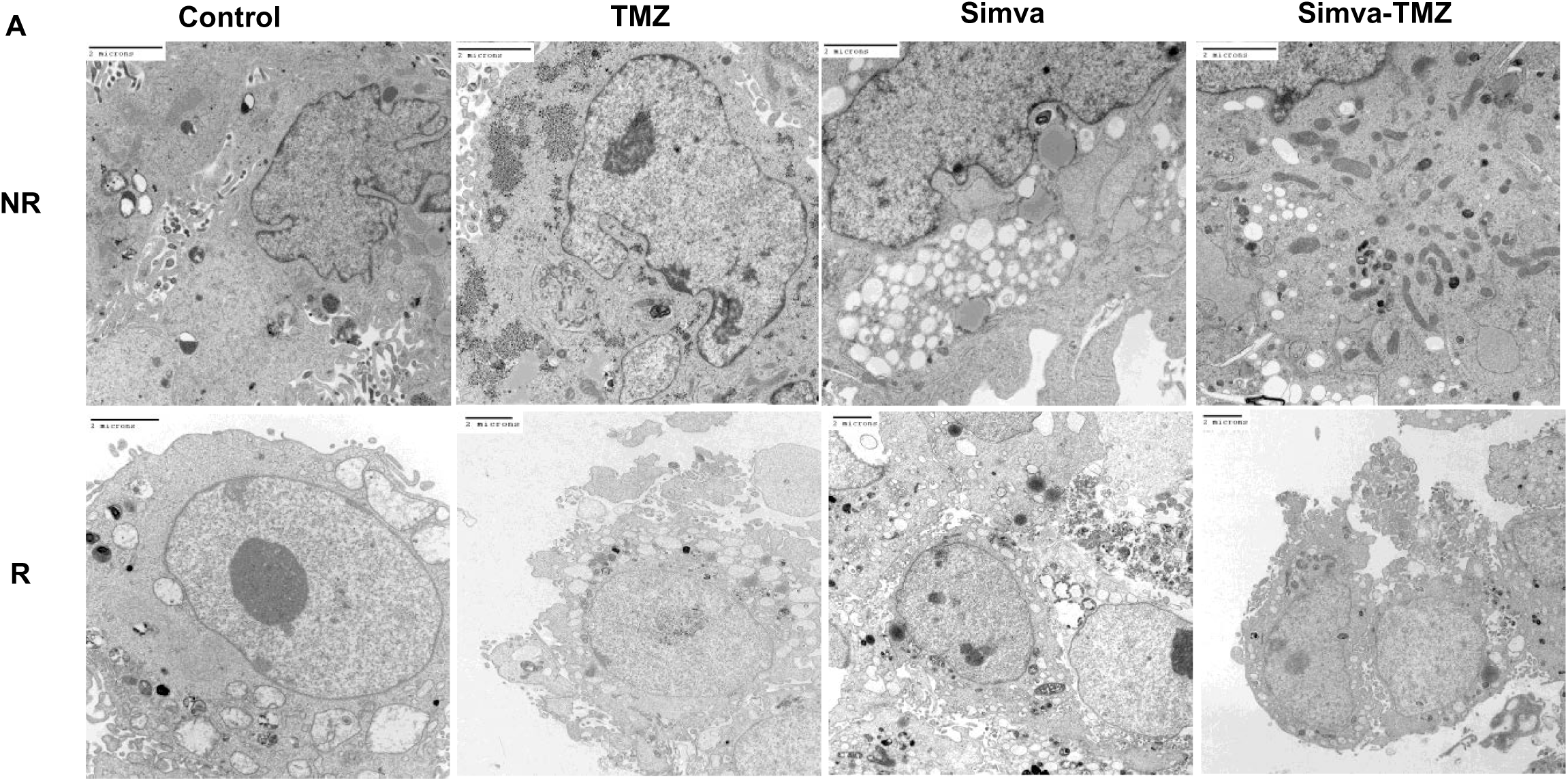
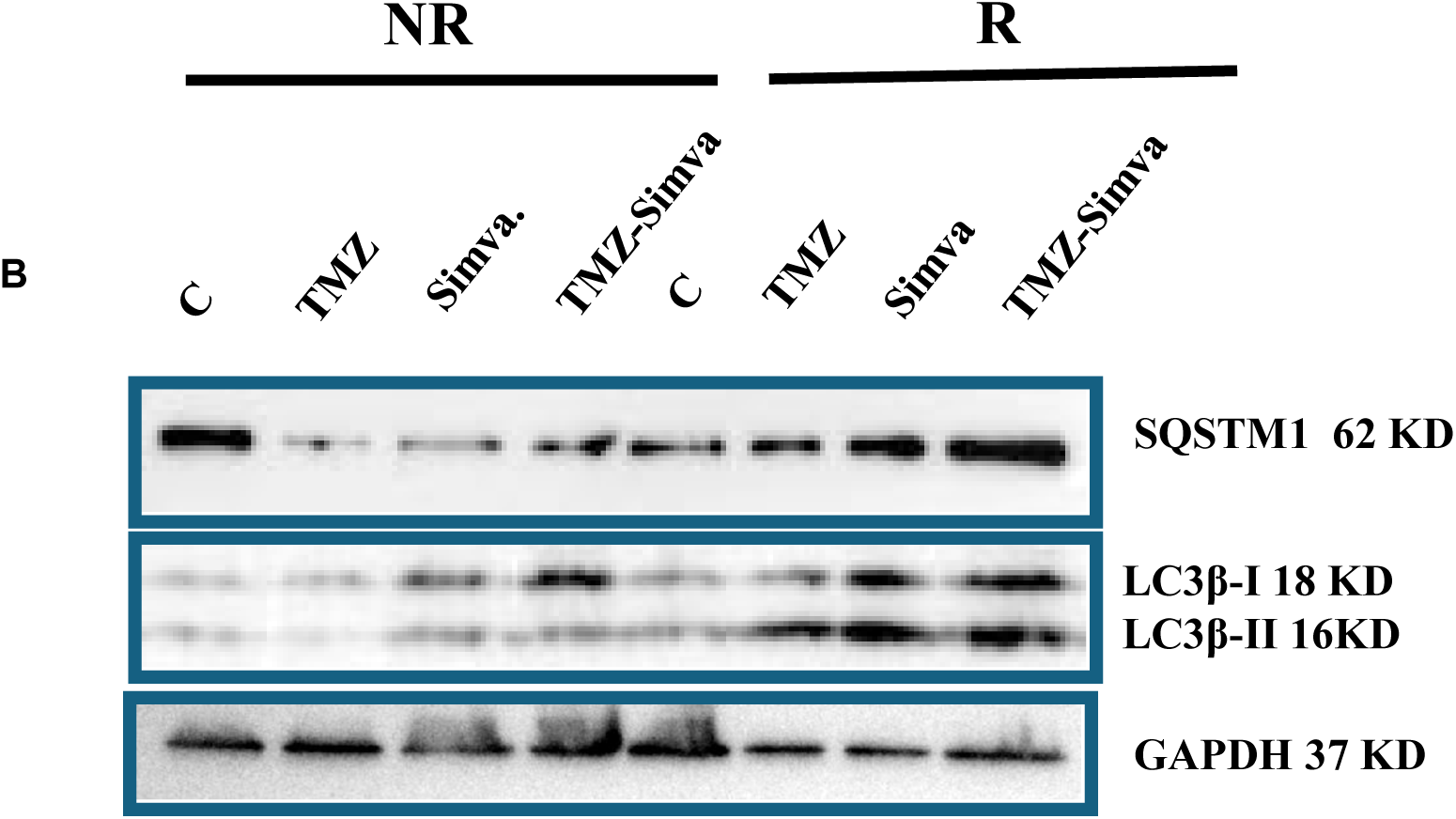
**TMZ, Simva and TMZ-Simva treatment do not change autophagy flux inhibition in U251 R cells**. U251 NR and R cells were treated with TMZ (100 µM), Simva (1 µ) and TMZ-Simva for 72 h. Autophagy flux was evaluated using TEM (**A**) and immunoblotting (**B**). TEM showed no further changes in number of double membrane vacuoles in R cells in all treatments while increased them in NR cells (**A**). Immunoblotting analysis showed that Simva and Simva-TMZ treatment induced LC3β-II accumulation and inhibits p62 degradation in NR cells while it did induce any significant changes in LC3β-II accumulation and inhibits p62 degradation in R cells (**B**). All experiments were done in 3 independent biological replicates ** means *P* ≤ 0.01, *** means *P* ≤ 0.001.

TEM analysis revealed no significant changes in the number or morphology of double-membrane autophagosomes in R cells across all treatment conditions, indicating persistent autophagosome accumulation irrespective of TMZ, Simva, or combination treatment (**Figure 6A**). In contrast, NR cells displayed a clear increase in autophagosome number following TMZ, Simva, or TMZ–Simva exposure, consistent with treatment-induced perturbation of autophagy dynamics.

Immunoblot analysis corroborated these ultrastructural findings. In NR cells, Simva and TMZ–Simva treatment resulted in accumulation of LC3β-II and impaired degradation of p62, indicative of autophagy flux inhibition (**Figure 6B**). By contrast, LC3β-II levels and p62 abundance remained unchanged in R cells following all treatments, consistent with an already established blockade of autophagy flux that is refractory to further pharmacologic modulation.

Together, these data demonstrate that neither TMZ nor simvastatin, alone or in combination, is sufficient to restore autophagy flux or alter autophagosome turnover in TMZ-resistant glioblastoma cells. This fixed autophagy-blocked state likely contributes to the stability of the resistant phenotype and underscores the challenge of targeting autophagy as a therapeutic vulnerability in established TMZ resistance.

## Discussion

Our findings reveal distinct autophagy marker patterns (LC3β and p62) in brain tumors and adjacent normal tissues, highlighting differences in autophagic activity. Tumor tissues exhibit increased autophagosome formation but impaired autophagic flux, leading to vesicle accumulation rather than degradation. In contrast, normal tissues show higher autophagy flux, maintaining metabolic balance. This dysregulated autophagy in tumors may sustain metabolic intermediates and enhance tumor survival under stress [63].

Autophagy and cholesterol biosynthesis play key roles in astrocytoma and GB pathogenesis. A strong correlation between LC3β puncta and autophagy activity suggests increased autophagosome formation as an adaptive mechanism to sustain energy and manage tumor metabolic stress [64, 65]. A positive correlation between LC3β puncta and UBIAD expression suggests autophagy supports cholesterol biosynthesis for tumor survival. Meanwhile, p62 accumulation inversely correlates with autophagy flux, indicating impaired autophagic degradation. In GB, UBIAD downregulation further links autophagy dysfunction to cholesterol metabolism dysregulation [66–68]. An important aspect highlighted by our tissue-based analyses is that dysregulation of autophagy and cholesterol metabolism is not a binary feature of glioblastoma but instead emerges progressively during glioma evolution from astrocytoma to glioblastoma. Astrocytomas already display elevated autophagosome accumulation and increased FDPS expression compared with normal brain tissue, indicating that autophagy–lipid coupling is engaged early in tumorigenesis. However, this state remains heterogeneous, with partial preservation of autophagic flux and metabolic flexibility. In contrast, glioblastomas exhibit a uniform phenotype characterized by robust autophagosome accumulation, near-complete loss of effective autophagic flux, and pronounced upregulation of FDPS, reflecting a fully established autophagy–cholesterol rewiring program. This stepwise intensification suggests that progressive uncoupling of autophagosome formation from degradation, together with reinforcement of cholesterol biosynthetic and buffering pathways, contributes to malignant progression and metabolic hardening. Thus, autophagy flux blockade and cholesterol pathway remodeling appear to function as adaptive layers that accumulate with tumor grade, predisposing high-grade gliomas to stress tolerance and therapeutic resistance. Thus, autophagy flux blockade and cholesterol pathway remodeling appear to function as adaptive layers that accumulate with tumor grade, predisposing high-grade gliomas to stress tolerance and therapeutic resistance in part through metabolic reprogramming and lipid homeostasis alterations [23, 69, 70].

TMZ-resistant GB cells undergo EMT, mitotic quiescence, and a metabolic shift to OXPHOS, enhancing survival. Resistant cells exhibit elongated morphology, increased vesicles, and upregulated EMT markers (N-cadherin, Vimentin), promoting invasiveness, plasticity, and therapy resistance [71, 72]. Blocking autophagy flux with Bafilomycin A1 did not affect EMT marker expression, suggesting EMT persists independently of autophagy. Resistant cells exhibit mitotic quiescence, marked by reduced proliferation, fewer mitotic cells, and G1 phase arrest, mirroring stem-like traits that enhance survival under chemotherapy [73]. Mitotic quiescence shields cells from TMZ-induced apoptosis, as reflected by a lower sub-G0-G1 population, and allows them to conserve energy and evade cellular stress associated with rapid proliferation [74]. This quiescent state, often linked to tumor-initiating cells, could serve as a reservoir for recurrence post-treatment, highlighting a critical therapeutic challenge in GB management [75].

The metabolic phenotype associated with temozolomide resistance in glioblastoma appears to reflect an adaptive state geared toward stress tolerance rather than enhanced bioenergetic output. Rather than increasing mitochondrial ATP production or respiratory reserve, resistant cells adopt a constrained energetic program characterized by reduced metabolic flexibility. This shift is consistent with a broader therapy-tolerant phenotype in which survival is favored over proliferation, aligning with the observed mitotic quiescence and reduced growth capacity of resistant cells. Elevated proton leak in resistant cells suggests adaptive mitochondrial uncoupling, a mechanism that may limit oxidative stress and preserve cellular integrity under chronic therapeutic pressure [76, 77]. Such uncoupling has been proposed as a survival strategy in cancer cells exposed to sustained stress, allowing maintenance of redox balance and mitochondrial function without committing to high-energy-demand states [77]. In this context, mitochondrial remodeling reflects functional adaptation rather than optimization of oxidative metabolism.

Additionally, our findings corroborate previous evidence [3], indicating that TMZ resistance is associated with inhibited autophagy flux. Autophagy can promote both survival and cell death, but in resistant cells, blocking autophagy flux allows vesicle accumulation, buffering metabolic stress and sustaining homeostasis. This aligns with studies showing autophagy flux inhibition as a strategy for cancer cells to evade apoptosis and adapt to chemotherapy [78].

In TMZ-resistant cells, we observed a blockade of autophagy flux, as indicated by autophagosome accumulation, disrupting lipophagy, the process responsible for degrading lipid droplets and recycling cholesterol. Cholesterol plays a critical role in regulating autophagy, particularly in the formation and maturation of autophagosomes. Cholesterol is an essential component of cellular membranes, including the membranes of autophagosomes and lysosomes, which are crucial for maintaining autophagy flux[62, 68, 79]. This impaired autophagic clearance reduces cholesterol availability, reflected by decreased de novo cholesterol synthesis and the downregulation of cholesterol biosynthesis genes, such as SREBP-2 and LDL-R. To compensate, R cells reprogram their lipid metabolism, shifting towards cholesterol esterification, as evidenced by the increase in specific cholesterol esters (CEs), including CE 22:5, CE 22:6, and CE 20:4. This metabolic shift allows R cells to store cholesterol in a non-toxic form, buffering against oxidative and metabolic stress while creating a reservoir of lipids that may support membrane synthesis and structural integrity under stress[16, 23, 80]. Despite clear evidence that temozolomide-resistant cells exhibit suppressed de novo cholesterol biosynthesis and transcriptional downregulation of LDL-R and SREBP-2, exogenous cholesterol delivery via lipid nanoparticles (LNPs) failed to restore chemosensitivity or reverse resistance. This finding suggests that cholesterol limitation per se is not the dominant vulnerability in resistant cells. Instead, resistant glioblastoma cells appear metabolically entrenched in a state that favors cholesterol sequestration and esterification rather than utilization for membrane synthesis or signaling. The observed lipotoxicity induced by LNPs in both resistant and non-resistant cells further supports the notion that excess free cholesterol cannot be productively integrated into cellular metabolism when autophagic lipid recycling is impaired. In resistant cells, autophagy flux blockade likely prevents efficient redistribution and turnover of exogenous lipids, rendering cholesterol supplementation ineffective as a resensitization strategy. These results emphasize that metabolic dependencies in chemoresistant glioblastoma are shaped not only by substrate availability but also by intracellular trafficking and degradation capacity. Therapeutic strategies aimed at overcoming resistance will therefore need to disrupt both lipid storage pathways and the autophagy machinery that buffers metabolic stress, rather than attempting to replenish depleted metabolites alone [23, 81, 82].

The ST and TMZ-ST treatments further highlight metabolic reprogramming. While TMZ alone induces minimal changes in CEs, the combined TMZ-ST treatment amplifies CE alterations, suggesting that targeting cholesterol metabolism can exacerbate stress in resistant cells [83–85]. GB relies on cholesterol esterification, driven by the increased activity of Sterol O-acyltransferase 1 (SOAT1), also known as acyl-CoA:cholesterol acyltransferase 1 (ACAT1), which converts free cholesterol into cholesteryl esters stored in lipid droplets. SOAT1 is overexpressed in GB, promoting lipid accumulation, tumor progression, and poor patient survival. Inhibiting SOAT1 reduces cholesterol esterification, leading to free cholesterol buildup in the endoplasmic reticulum (ER). This disrupts lipid metabolism by suppressing sterol regulatory element-binding protein-1 (SREBP-1), reducing fatty acid synthesis and impairing tumor growth. Preclinical studies show that genetic silencing of SOAT1 and pharmacological inhibition (e.g., avasimibe and K604) significantly decrease GB cell viability and tumor progression [23, 86–89]. Impaired autophagy flux further exacerbates this dependency by limiting the clearance of lipid droplets and organelles, reinforcing the need for alternative lipid storage mechanisms. This interplay between autophagy and lipid metabolism represents a dual vulnerability in R cells. The blockade of autophagy flux increases dependency on CE accumulation and disrupts cellular homeostasis, forcing metabolic adaptations to sustain chemoresistance. The accumulation of cholesteryl esters and triglycerides in high-grade tumor extracts is associated with the extent of vascular proliferation, distinguishing them from both normal brain tissue and low-grade neoplasms[90].

Our findings reveal a compelling connection between autophagy dysregulation and metabolic reprogramming in R GB cells, providing key insights into mechanisms of chemoresistance. KEGG pathway enrichment identified Steroid hormone biosynthesis, Ovarian steroidogenesis, Neuroactive ligand-receptor interaction, Insulin resistance, and Metabolic pathways as the top pathways differentiating R and TMZ-sensitive (NR) cells. These pathways are intricately linked to autophagy, underscoring its central role in regulating cellular adaptation. Steroid hormones, including glucocorticoids, are known to modulate autophagy through signaling pathways that converge on mTOR-dependent and stress-responsive regulatory networks, with mTOR acting as a central integrator of hormonal and metabolic cues controlling autophagic activity [91, 92]. Similarly, steroid hormones associated with ovarian steroidogenesis, including estrogen and progesterone, have been reported to modulate autophagy in specific cancer contexts [93–95]. The Neuroactive ligand-receptor interaction pathway highlights the role of neurotransmitters and calcium signaling in regulating autophagy [96, 97], while Insulin resistance disrupts glucose metabolism and activates stress responses that directly alter autophagic flux [98]. Finally, Metabolic pathways integrate lipid and energy metabolism with autophagic processes, coordinating lipid turnover and homeostasis [29, 99, 100].

This study demonstrates that glioblastoma progression and temozolomide resistance are underpinned by a coordinated interconnection of autophagy and lipid metabolism rather than by isolated pathway alterations. Brain tumors exhibit a characteristic pattern of increased autophagosome formation coupled with impaired autophagic flux, distinguishing malignant tissue from adjacent normal brain and establishing vesicle accumulation as a hallmark of tumor-associated metabolic stress adaptation. In glioblastoma, this dysregulated autophagy state is tightly linked to alterations in cholesterol handling, including suppressed de novo cholesterol synthesis, transcriptional downregulation of SREBP-2 and LDL-R, and compensatory cholesterol ester accumulation.

In the context of temozolomide resistance, these alterations converge into a stable, therapy-tolerant phenotype characterized by epithelial-to-mesenchymal transition, mitotic quiescence, constrained bioenergetic capacity, and adaptive mitochondrial uncoupling. Rather than enhancing ATP production, resistant cells adopt an energy-conserving state that limits oxidative stress and preserves cellular integrity under chronic therapeutic pressure. Importantly, pharmacologic modulation of cholesterol biosynthesis or autophagy flux alone is insufficient to restore chemosensitivity, underscoring the robustness of this adaptive program.

Together, our findings identify autophagy flux blockade and cholesterol esterification as interconnected features of glioblastoma resistance biology. This autophagy–lipid axis supports survival under treatment stress and represents a central organizing principle of chemoresistance, providing a unifying framework that links vesicle dynamics, metabolic plasticity, and therapeutic failure in glioblastoma.

These results highlight several important avenues for future investigation. First, mechanistic studies are needed to define how autophagy flux blockade is established and maintained in resistant glioblastoma cells, including the upstream regulatory events that decouple autophagosome formation from lysosomal degradation. Dissecting whether this blockade arises from lysosomal dysfunction, membrane lipid composition changes, or signaling rewiring will be critical.

Second, the reliance of resistant cells on cholesterol esterification suggests that simultaneous targeting of lipid storage pathways (e.g., SOAT1/ACAT1 inhibition) and autophagy regulation may represent a more effective strategy than single-pathway interventions. Future work should evaluate combinatorial approaches that disrupt both lipid buffering capacity and vesicle turnover to destabilize the resistant metabolic state.

Third, the metabolic adaptations observed here—mitotic quiescence, reduced respiratory reserve, and adaptive mitochondrial uncoupling—raise the possibility that resistance is sustained by stress-tolerant, low-proliferative cell populations. Longitudinal and single-cell analyses will be essential to determine whether these cells serve as reservoirs for recurrence and how they transition back to proliferative states following treatment withdrawal.

Finally, extending these findings to patient-derived models and clinical samples will be crucial to establish the translational relevance of the autophagy–cholesterol axis. Biomarkers reflecting autophagy flux status and lipid metabolic remodeling may aid in stratifying patients and guiding rational combination therapies aimed at overcoming temozolomide resistance.

## Author Contirbution

**Shahla Shojaei** and **Amir Barzegar Behrooz** contributed to drafting the manuscript and performing experiments; **Kianoosh Naghibzadeh**, **João Basso**, **Tania Dehesh**, **Chris Pasco**, **Amir Ravandi**, **Rui Vitorino**, and **Stevan Pešić** performed lipidomics and related analyses; **Javad Alizadeh** conducted autophagy assays; **Roham Sabei** and **Bhavya Bhushan** performed experiments related to Figure 6; **Mehdi Eshraghi** contributed to manuscript drafting; **Simone De Silva Rosa**, **Courtney Clark**, **Vinith Yathindranath**, and **Donald Miller** performed and led LNP-related studies; **Mateusz Tomczyk**, **Laura Cole**, **Grant Hatch**, and **Vern Dolinsky** conducted LDL, SREBP, and cholesterol metabolism studies; **Sanjiv Dhingra** and **Abhay Srivastava** performed mitochondrial metabolism studies; **Negar Azarpira** conducted tissue pathology analysis; **Mahmoud Aghaei** performed tissue microarray and correlation analyses; **Saeid Ghavami** conceived and led the project, supervised the research, coordinated collaborations, and finalized the manuscript.

## Acknowledgment

We acknowledge the use of ChatGPT (version 5.2, OpenAI) for supervised grammar editing and language refinement of the manuscript.

## References

1. Emir, S.M., et al., Hunting glioblastoma recurrence: glioma stem cells as retrospective targets. Am J Physiol Cell Physiol, 2025. 328(3): p. C1045–c1061.

2. Pouyan, A., et al., Glioblastoma multiforme: insights into pathogenesis, key signaling pathways, and therapeutic strategies. Mol Cancer, 2025. 24(1): p. 58.

3. Yan, Y., et al., Targeting autophagy to sensitive glioma to temozolomide treatment. J Exp Clin Cancer Res, 2016. 35: p. 23.

4. Teraiya, M., H. Perreault, and V.C. Chen, An overview of glioblastoma multiforme and temozolomide resistance: can LC-MS-based proteomics reveal the fundamental mechanism of temozolomide resistance? Front Oncol, 2023. 13: p. 1166207.

5. Wu, H., et al., Research progress of drug resistance mechanism of temozolomide in the treatment of glioblastoma. Heliyon, 2024. 10(21): p. e39984.

6. Hwang, Y.K., et al., Importance of Autophagy Regulation in Glioblastoma with Temozolomide Resistance. Cells, 2024. 13(16).

7. Koh, M., et al., ANXA2 (annexin A2) is crucial to ATG7-mediated autophagy, leading to tumor aggressiveness in triple-negative breast cancer cells. Autophagy, 2024. 20(3): p. 659–674.

8. Wang, H., et al., Long Non-Coding RNA NNT-AS1 Contributes to Cisplatin Resistance via miR-1236-3p/ATG7 Axis in Lung Cancer Cells. Onco Targets Ther, 2020. 13: p. 3641–3652.

9. Yu, W., L. Ma, and X. Li, DANCR promotes glioma cell autophagy and proliferation via the miR-33b/DLX6/ATG7 axis. Oncol Rep, 2023. 49(2).

10. Qin, Y., et al., Autophagy machinery in glioblastoma: The prospect of cell death crosstalk and drug resistance with bioinformatics analysis. Cancer Letters, 2024. 580: p. 216482.

11. Clark, C., et al., Assessing Autophagy Flux in Glioblastoma Temozolomide Resistant Cells. Methods Mol Biol, 2025. 2879: p. 225–238.

12. Clark, C., et al., *BCL2L13 Influences Autophagy and Ceramide Metabolism without Affecting Temozolomide Resistance in Glioblastoma.* bioRxiv, 2024.

13. Meena, D. and S. Jha, Autophagy in glioblastoma: A mechanistic perspective. International Journal of Cancer, 2024. 155(4): p. 605–617.

14. DeBerardinis, R.J., et al., The biology of cancer: metabolic reprogramming fuels cell growth and proliferation. Cell Metab, 2008. 7(1): p. 11–20.

15. Eyme, K.M., et al., Targeting de novo lipid synthesis induces lipotoxicity and impairs DNA damage repair in glioblastoma mouse models. Sci Transl Med, 2023. 15(679): p. eabq6288.

16. Weng, X., et al., Lipidomics-driven drug discovery and delivery strategies in glioblastoma. Biochim Biophys Acta Mol Basis Dis, 2025. 1871(3): p. 167637.

17. Kao, T.J., et al., Dysregulated lipid metabolism in TMZ-resistant glioblastoma: pathways, proteins, metabolites and therapeutic opportunities. Lipids Health Dis, 2023. 22(1): p. 114.

18. Yamamoto, Y., et al., Intracellular cholesterol level regulates sensitivity of glioblastoma cells against temozolomide-induced cell death by modulation of caspase-8 activation via death receptor 5-accumulation and activation in the plasma membrane lipid raft. Biochem Biophys Res Commun, 2018. 495(1): p. 1292–1299.

19. Ahmad, F., et al., Cholesterol Metabolism: A Potential Therapeutic Target in Glioblastoma. Cancers (Basel), 2019. 11(2).

20. Villa, G.R., et al., An LXR-Cholesterol Axis Creates a Metabolic Co-Dependency for Brain Cancers. Cancer Cell, 2016. 30(5): p. 683–693.

21. Sato, R., Sterol metabolism and SREBP activation. Arch Biochem Biophys, 2010. 501(2): p. 177–81.

22. Yahagi, N., et al., A crucial role of sterol regulatory element-binding protein-1 in the regulation of lipogenic gene expression by polyunsaturated fatty acids. J Biol Chem, 1999. 274(50): p. 35840–4.

23. Guo, X., et al., Cholesterol metabolism and its implication in glioblastoma therapy. J Cancer, 2022. 13(6): p. 1745–1757.

24. Meyer, N., et al., Autophagy activation, lipotoxicity and lysosomal membrane permeabilization synergize to promote pimozide- and loperamide-induced glioma cell death. Autophagy, 2021. 17(11): p. 3424–3443.

25. Geng, F., et al., SREBP-1 upregulates lipophagy to maintain cholesterol homeostasis in brain tumor cells. Cell Reports, 2023. 42(7): p. 112790.

26. Kanzawa, T., et al., Role of autophagy in temozolomide-induced cytotoxicity for malignant glioma cells. Cell Death Differ, 2004. 11(4): p. 448–57.

27. Xia, Q., et al., Therapeutic Potential of Autophagy in Glioblastoma Treatment With Phosphoinositide 3-Kinase/Protein Kinase B/Mammalian Target of Rapamycin Signaling Pathway Inhibitors. Front Oncol, 2020. 10: p. 572904.

28. Hu, Y.L., et al., Hypoxia-induced autophagy promotes tumor cell survival and adaptation to antiangiogenic treatment in glioblastoma. Cancer Res, 2012. 72(7): p. 1773–83.

29. Xie, Y., et al., Interplay Between Lipid Metabolism and Autophagy. Front Cell Dev Biol, 2020. 8: p. 431.

30. Zhang, S., et al., The regulation, function, and role of lipophagy, a form of selective autophagy, in metabolic disorders. Cell Death Dis, 2022. 13(2): p. 132.

31. Li, X., et al., Crosstalk between lipid droplets and autophagy in cancer: A nexus for therapeutic targeting. Pharmacol Res, 2025. 222: p. 108023.

32. McDermott, M., et al., In vitro Development of Chemotherapy and Targeted Therapy Drug-Resistant Cancer Cell Lines: A Practical Guide with Case Studies. Front Oncol, 2014. 4: p. 40.

33. Adlimoghaddam, A., et al., Sex and region-specific disruption of autophagy and mitophagy in Alzheimer’s disease: linking cellular dysfunction to cognitive decline. Cell Death Discov, 2025. 11(1): p. 204.

34. Alizadeh, J., et al., Evaluation of Mitochondrial Phagy (Mitophagy) in Human Non-small Adenocarcinoma Tumor Cells. Methods Mol Biol, 2025. 2879: p. 261–273.

35. Alizadeh, J., et al., Mevalonate Cascade Inhibition by Simvastatin Induces the Intrinsic Apoptosis Pathway via Depletion of Isoprenoids in Tumor Cells. Sci Rep, 2017. 7: p. 44841.

36. Shojaei, S., et al., Simvastatin increases temozolomide-induced cell death by targeting the fusion of autophagosomes and lysosomes. FEBS J, 2020. 287(5): p. 1005–1034.

37. Ghavami, S., et al., Statin-triggered cell death in primary human lung mesenchymal cells involves p53-PUMA and release of Smac and Omi but not cytochrome c. Biochim Biophys Acta, 2010. 1803(4): p. 452–67.

38. Folch, J., M. Lees, and G.H. Sloane Stanley, A simple method for the isolation and purification of total lipides from animal tissues. J Biol Chem, 1957. 226(1): p. 497–509.

39. Cole, L.K., R.L. Jacobs, and D.E. Vance, Tamoxifen induces triacylglycerol accumulation in the mouse liver by activation of fatty acid synthesis. Hepatology, 2010. 52(4): p. 1258–65.

40. Basso, J., et al., Are we better together? Addressing a combined treatment of pitavastatin and temozolomide for brain cancer. Eur J Pharmacol, 2024. 985: p. 177087.

41. Pirmoradi, L., et al., Targeting cholesterol metabolism in glioblastoma: a new therapeutic approach in cancer therapy. J Investig Med, 2019. 67(4): p. 715–719.

42. Shojaei, S., et al., *Unlocking a New Path: An Autophagometer that Measures Flux Using a Non-Fluorescent Immunohistochemistry Method*. bioRxiv, 2024.

43. Ershov, P., et al., Enzymes in the Cholesterol Synthesis Pathway: Interactomics in the Cancer Context. Biomedicines, 2021. 9(8).

44. Juarez, D. and D.A. Fruman, Targeting the Mevalonate Pathway in Cancer. Trends Cancer, 2021. 7(6): p. 525–540.

45. Sethunath, V., et al., Targeting the Mevalonate Pathway to Overcome Acquired Anti-HER2 Treatment Resistance in Breast Cancer. Mol Cancer Res, 2019. 17(11): p. 2318–2330.

46. Ghavami, S., et al., Mevalonate cascade regulation of airway mesenchymal cell autophagy and apoptosis: a dual role for p53. PLoS One, 2011. 6(1): p. e16523.

47. Toepfer, N., et al., Atorvastatin induces autophagy in prostate cancer PC3 cells through activation of LC3 transcription. Cancer Biol Ther, 2011. 12(8): p. 691–9.

48. Yang, Z., et al., Fluvastatin Prevents Lung Adenocarcinoma Bone Metastasis by Triggering Autophagy. EBioMedicine, 2017. 19: p. 49–59.

49. Bernhard, C., et al., *Glioblastoma Metabolism: Insights and Therapeutic Strategies.* Int J Mol Sci, 2023. 24(11).

50. Davodabadi, F., et al., Cancer chemotherapy resistance: Mechanisms and recent breakthrough in targeted drug delivery. Eur J Pharmacol, 2023. 958: p. 176013.

51. Singh, R. and A.M. Cuervo, Autophagy in the cellular energetic balance. Cell Metab, 2011. 13(5): p. 495–504.

52. Dekker, L.J.M., et al., Multiomics profiling of paired primary and recurrent glioblastoma patient tissues. Neurooncol Adv, 2020. 2(1): p. vdaa083.

53. Liu, C., et al., New insights into the therapeutic potentials of statins in cancer. Front Pharmacol, 2023. 14: p. 1188926.

54. Preta, G., New Insights Into Targeting Membrane Lipids for Cancer Therapy. Front Cell Dev Biol, 2020. 8: p. 571237.

55. 55. Hsu, T.I., Editorial: Metabolic reprogramming for acquiring therapeutic resistance in glioblastoma. Front Oncol, 2023. 13: p. 1220063.

56. Philipsen, M.H., et al., Distinct Cholesterol Localization in Glioblastoma Multiforme Revealed by Mass Spectrometry Imaging. ACS Chem Neurosci, 2023. 14(9): p. 1602–1609.

57. Shen, G., et al., Autophagy as a target for glucocorticoid-induced osteoporosis therapy. Cell Mol Life Sci, 2018. 75(15): p. 2683–2693.

58. Afzal, A., et al., Functional role of autophagy in testicular and ovarian steroidogenesis. Front Cell Dev Biol, 2024. 12: p. 1384047.

59. Ho, S.M., Estrogen, progesterone and epithelial ovarian cancer. Reprod Biol Endocrinol, 2003. 1: p. 73.

60. Menzies, F.M., et al., *Autophagy and Neurodegeneration: Pathogenic Mechanisms and Therapeutic Opportunities.* Neuron, 2017. 93(5): p. 1015–1034.

61. Sadeghi, A., et al., Crosstalk between autophagy and insulin resistance: evidence from different tissues. Eur J Med Res, 2023. 28(1): p. 456.

62. Jarocki, M., et al., Lipids associated with autophagy: mechanisms and therapeutic targets. Cell Death Discov, 2024. 10(1): p. 460.

63. Goldsmith, J., B. Levine, and J. Debnath, *Autophagy and cancer metabolism.* Methods Enzymol, 2014. 542: p. 25–57.

64. Russell, R.C. and K.L. Guan, The multifaceted role of autophagy in cancer. The EMBO Journal, 2022. 41(13): p. e110031.

65. Debnath, J., N. Gammoh, and K.M. Ryan, Autophagy and autophagy-related pathways in cancer. Nat Rev Mol Cell Biol, 2023. 24(8): p. 560–575.

66. Xiao, M., et al., Functional significance of cholesterol metabolism in cancer: from threat to treatment. Exp Mol Med, 2023. 55(9): p. 1982–1995.

67. Xia, W., et al., The role of cholesterol metabolism in tumor therapy, from bench to bed. Front Pharmacol, 2023. 14: p. 928821.

68. Shapira, K.E., et al., Autophagy is induced and modulated by cholesterol depletion through transcription of autophagy-related genes and attenuation of flux. Cell Death Discov, 2021. 7(1): p. 320.

69. Abdul Rashid, K., et al., Lipid Alterations in Glioma: A Systematic Review. Metabolites, 2022. 12(12).

70. Escamilla-Ramirez, A., et al., Autophagy as a Potential Therapy for Malignant Glioma. Pharmaceuticals (Basel), 2020. 13(7).

71. Thiery, J.P., et al., Epithelial-mesenchymal transitions in development and disease. Cell, 2009. 139(5): p. 871–90.

72. Tang, H., et al., SRPX2 Enhances the Epithelial-Mesenchymal Transition and Temozolomide Resistance in Glioblastoma Cells. Cell Mol Neurobiol, 2016. 36(7): p. 1067–76.

73. Zhao, J., Cancer stem cells and chemoresistance: The smartest survives the raid. Pharmacol Ther, 2016. 160: p. 145–58.

74. Lindell, E., L. Zhong, and X. Zhang, Quiescent Cancer Cells—A Potential Therapeutic Target to Overcome Tumor Resistance and Relapse. International Journal of Molecular Sciences, 2023. 24(4): p. 3762.

75. Antonica, F., et al., A slow-cycling/quiescent cells subpopulation is involved in glioma invasiveness. Nat Commun, 2022. 13(1): p. 4767.

76. Brand, M.D. and D.G. Nicholls, Assessing mitochondrial dysfunction in cells. Biochem J, 2011. 435(2): p. 297–312.

77. Divakaruni, A.S. and M.D. Brand, The regulation and physiology of mitochondrial proton leak. Physiology (Bethesda), 2011. 26(3): p. 192–205.

78. Kimmelman, A.C., The dynamic nature of autophagy in cancer. Genes Dev, 2011. 25(19): p. 1999–2010.

79. Maharjan, Y., et al., Intracellular cholesterol transport inhibition Impairs autophagy flux by decreasing autophagosome-lysosome fusion. Cell Commun Signal, 2022. 20(1): p. 189.

80. Choo, M., et al., Involvement of cell shape and lipid metabolism in glioblastoma resistance to temozolomide. Acta Pharmacol Sin, 2023. 44(3): p. 670–679.

81. Tonkin-Reeves, A., C.M. Giuliani, and J.T. Price, Inhibition of autophagy; an opportunity for the treatment of cancer resistance. Front Cell Dev Biol, 2023. 11: p. 1177440.

82. Jusovic, M., et al., The Combined Inhibition of Autophagy and Diacylglycerol Acyltransferase - Mediated Lipid Droplet Biogenesis Induces Cancer Cell Death during Acute Amino Acid Starvation. Cancers (Basel), 2023. 15(19).

83. Yamamoto, Y., et al., Involvement of Intracellular Cholesterol in Temozolomide-Induced Glioblastoma Cell Death. Neurol Med Chir (Tokyo), 2018. 58(7): p. 296–302.

84. Alrosan, A.Z., et al., The effects of statin therapy on brain tumors, particularly glioma: a review. Anticancer Drugs, 2023. 34(9): p. 985–994.

85. Wu, H., et al., Effect of simvastatin on glioma cell proliferation, migration, and apoptosis. Neurosurgery, 2009. 65(6): p. 1087–96; discussion 1096-7.

86. Geng, F., et al., Inhibition of SOAT1 Suppresses Glioblastoma Growth via Blocking SREBP-1-Mediated Lipogenesis. Clin Cancer Res, 2016. 22(21): p. 5337–5348.

87. Bemlih, S., M.D. Poirier, and A. El Andaloussi, Acyl-coenzyme A: cholesterol acyltransferase inhibitor Avasimibe affect survival and proliferation of glioma tumor cell lines. Cancer Biol Ther, 2010. 9(12): p. 1025–32.

88. Sun, T. and X. Xiao, Targeting ACAT1 in cancer: from threat to treatment. Frontiers in Oncology, 2024. 14.

89. Ohmoto, T., et al., K604, a specific acyl-CoA:cholesterol acyltransferase 1 inhibitor, suppresses proliferation of U251-MG glioblastoma cells. Mol Med Rep, 2015. 12(4): p. 6037–42.

90. Tugnoli, V., et al., Characterization of lipids from human brain tissues by multinuclear magnetic resonance spectroscopy. Biopolymers, 2001. 62(6): p. 297–306.

91. Galluzzi, L. and D.R. Green, Autophagy-Independent Functions of the Autophagy Machinery. Cell, 2019. 177(7): p. 1682–1699.

92. Laplante, M. and D.M. Sabatini, mTOR signaling in growth control and disease. Cell, 2012. 149(2): p. 274–93.

93. Cook, K.L. and R. Clarke, Estrogen receptor-alpha signaling and localization regulates autophagy and unfolded protein response activation in ER+ breast cancer. Receptors Clin Investig, 2014. 1(6).

94. Raut, P.K., et al., Estrogen receptor signaling mediates leptin-induced growth of breast cancer cells via autophagy induction. Oncotarget, 2017. 8(65): p. 109417–109435.

95. Sui, X., et al., Autophagy and chemotherapy resistance: a promising therapeutic target for cancer treatment. Cell Death Dis, 2013. 4(10): p. e838.

96. Sukumaran, P., et al., Calcium Signaling Regulates Autophagy and Apoptosis. Cells, 2021. 10(8).

97. Bootman, M.D., et al., The regulation of autophagy by calcium signals: Do we have a consensus? Cell Calcium, 2018. 70: p. 32–46.

98. Zhang, N., et al., Autophagy regulates insulin resistance following endoplasmic reticulum stress in diabetes. J Physiol Biochem, 2015. 71(2): p. 319–27.

99. Yang, Y., et al., Interplay of CD36, autophagy, and lipid metabolism: insights into cancer progression. Metabolism, 2024. 155: p. 155905.

100. Saito, T., et al., Autophagy regulates lipid metabolism through selective turnover of NCoR1. Nat Commun, 2019. 10(1): p. 1567.

